# Smoking, asthma and airway microbial disruption

**DOI:** 10.1101/583559

**Authors:** Elena M. Turek, Michael J. Cox, Michael Hunter, Jennie Hui, Phillip James, Saffron A.G. Willis-Owen, Leah Cuthbertson, Alan James, A. William Musk, Miriam F. Moffatt, William O.C.M. Cookson

## Abstract

**Background:** Normal airway microbial communities play a central role in respiratory health but are poorly characterized. Cigarette smoking is the dominant global environmental influence on lung function, and asthma has become the most prevalent chronic respiratory disease worldwide. Both conditions have major microbial components that are also poorly defined.

**Methods:** We investigated airway bacterial communities in a general population sample of 529 Australian adults. Posterior oropharyngeal swabs were analysed by sequencing of the 16S rRNA and methionine aminopeptidase genes. The microbiota were characterised according to their prevalence, abundance, and network memberships.

**Findings:** Microbial communities were similar across the population and were strongly organized into co-abundance networks. Smoking associated with diversity loss, negative effects on abundant taxa, profound alterations to network structure and expansion of *Streptococcus* spp. By contrast, the asthmatic microbiota were selectively affected by an increase in *Neisseria* spp. and by reduced numbers of low abundance but prevalent organisms.

**Interpretation:** Our study shows healthy airway microbiota are contained within a highly structured ecosystem, indicating balanced relationships between the microbiome and human host factors. The marked abnormalities in smokers may be pathogenic for chronic obstructive pulmonary disease (COPD) and lung cancer. The narrow spectrum of abnormalities in asthmatics encourages investigation of damaging and protective effects of specific bacteria.

**Funding:** The study was funded by the Asmarley Trust and a Wellcome Senior Investigator Award to WOCC and MFM (P46009). The Busselton Healthy Ageing Study is supported by the Government of Western Australia (Office of Science, Department of Health) the City of Busselton, and private donations.

## Introduction

All animals and plants establish symbiotic relationships with microorganisms^1^, to the extent that hosts and their associated microorganisms can be considered to be a co-evolving unit known as a “holobiont”^1^. The airways of the human lung carry commensal microbiota at a similar density to the small intestine^2^ and the surface area of the lungs is greater than that of the gut (40 to 80m^2^ compared to 30m^2^) so that the respiratory microbiota have profound opportunities to affect health^3,4^. The ecology of normal airway microbial communities and the means through which they modify disease are however still poorly understood, and there is as yet no systematic basis for their study.

A quarter of men and 5% of women in the world smoke cigarettes daily^5^. Smoking causes 11·5% of deaths globally^5^ and COPD and lung cancer are its most common pulmonary consequences. COPD is accompanied by recurrent infections^6,7^ and bacteria contribute to lung carcinogenesis^8^.

Asthma is an inflammatory disorder of the airways that has become the most prevalent chronic respiratory disease worldwide^9,10^. In numerous studies its rise has been linked to urbanization and the loss of traditional rural environments^11–13^. The “hygiene hypothesis” suggests that loss of microbial exposure allows asthma to develop^14,15^. Formulation and testing of potential mechanisms through which a rich microbial environment may protect against asthma has been restrained by limited understanding of the airway microbiota in epidemiological samples.

We therefore examined adults from the population of Busselton in Western Australia who were participating in a general health survey^16^. Direct sampling of the lung microbiota requires invasive procedures such as bronchoscopy that are not possible in epidemiological studies. In healthy individuals the microbiota of the oropharynx and the intra-thoracic airways are very similar^2,17^. We consequently used oropharyngeal swabs, taken beyond the tonsils and palate, for population sampling. We accept nevertheless that the abundance of pathogens in the lower airways of diseased subjects is only partially reflected in the oropharynx^2,18^.

We used PCR and sequencing of the 16S rRNA gene to identify bacterial taxa (operational taxonomic units, OTUs) present in the samples. However, *Streptococcus* spp., which are particularly prevalent in respiratory samples, show high rates of clonal diversity and are poorly differentiated by 16S sequences and by standard culture^19,20^. We therefore sequenced the methionine aminopeptidase gene *(map)* to further differentiate between *Streptococcus* taxa^20^.

Microbial communities are formed through complex ecological interactions that can be exposed through network analyses^21^. Assuming that correlations in the abundance of different taxa reflect co-ordinated growth, we applied weighted correlation network analyses (WGCNA)^22^ to the dataset. We aimed to detect major networks of interacting bacteria (“Guilds”) within airway communities, before testing how they were influenced by cigarette smoking or the presence of asthma.

## Methods

### Subject recruitment

Five hundred and 78 Caucasian adults were recruited through the Busselton Health Study in Western Australia. The study has received ethics approval from the University of Western Australia Human Research Ethics Committee (Number RA/4/1/2203). Individuals with a diagnosis of cancer were excluded. Subjects were not included if they were taking antibiotics within six weeks of the time of study. Participants completed a detailed questionnaire as previously described^16^. Subjects were classified as asthmatic if they answered yes to the question “Has your doctor ever told you that you have asthma”. Other diagnoses potentially influencing the microbiome were diabetes (n=18 patients) and gastro-esophageal reflux (GERD, n=36). No associations were found for diabetes or GERD in any analyses, and we included subjects with these diagnoses in the unaffected group.

Samples for microbial analysis were taken under direct vision, using sterile rayon swabs that were rubbed gently with an even pressure around the posterior oropharynx five times, strictly avoiding contact with tongue, tonsils, palate or nose. Swabs were immediately frozen and stored at −80°C prior to transportation on dry ice to Imperial College London, UK.

### 16S rRNA gene sequencing

DNA was extracted from swab heads using the MP Bio FastDNA Spin Kit for Soil (http://www.mpbio.com). A single sample was examined for each subject. Blank controls with no sample added were taken from each DNA extraction kit to test for contamination^23^.

PCR of the 16S rRNA V4 region was performed in quadruplicate using a custom indexed forward primer S-D-Bact-0564-a-S-15 (5’ AYT GGG YDT AAA GNG 3’), reverse primer S-D-Bact-0785-b-A-18 (5’ TAC NVG GGT ATC TAA TCC 3’) and a high fidelity Taq polymerase master mix (Q5, New England Biolabs, Massachusetts, USA). Primer sequences were based on Klindworth *et al*.^24^, with dual-barcoding as per Kozich *et al*.^25^ with adaptors from Illumina (California, USA). A mock community^26^ was included to assess sequencing quality. PCR cycling conditions were: 95oC for 2 minutes followed by 35 cycles of 95oC for 20 seconds, 50oC for 20 seconds and 72oC for 5 minutes. Amplicons were purified, quantified and equi-molar pooled and the library paired-end sequenced (Illumina MiSeq V2 reagent kit) as previously described^26^. Bacterial load was quantified by qPCR using KAPA BioSystems SYBR Fast qPCR Kit with the same 16S rRNA V4 primers used for sequencing.

Analysis of data was carried out in the R environment and details can be followed on github: https://tinyurl.com/y2onjblt. Sequence processing was performed in QIIME (Version 1.9.0)^27^. Community level differences in alpha and beta diversity and Operational Taxonomic Unit (OTU) level differences, were analysed using Phyloseq in R (Version 3.2.0). A phylogenetic tree was generated from the representative sequences using the default parameters of the make_phylogeny command^27^. Taxonomy of OTUs was assigned by matching representative sequences against release version 23 August 2013 of the Silva database^28^ using the default parameters of the assign_taxonomy command^27^. OTUs occurring in only one sample or with less than 20 reads in the whole dataset were removed. Weighted and unweighted UniFrac beta diversity measures and subsequent principal co-ordinates analysis of them was carried out using the beta_diversity_through_plots script^27^. For the purposes of alpha diversity calculations, the raw counts tables were rarefied to a minimum of 6,543 reads. Significant differences in alpha diversity between datasets were assessed using Mann–Whitney U-tests.

Potential kit contaminant OTUs were identified by the presence of negative Spearman’s correlations between OTU abundance and bacterial burden (logged qPCR copy number), adjusted using Bonferroni corrected P-values < 0.05. OTUs subsequently of interest were cross-checked with a listing of potential contaminants^23^.

### *Map* gene sequencing

We further differentiated *Streptococcus* spp. by sequencing the methionine aminopeptidase *(map)* gene^20^ in 483 samples (constrained to 5 sequencing runs with controls). Of these subjects 234 were never-smoking and 53 were current smokers. We used barcoded primers map-up 5’ GCW GACTCWT GTT GGGCWTAT GC ‘3 and map-down 5’ TTARTAAGTTCYTTCTTCDCCTTG ‘3. As positive controls, DNA from nine strains of *Streptococcus* with bacterial identity confirmed through Sanger sequencing was used for positive controls (S. agalactiae (DSMZ-2134); *S. constellatus* subsp. *Constellatus* (DSMZ-20575); *S. infantis* (DSMZ-12492); *S. parasanguinis* (DSMZ-6778); *S. pneumoniae* (DSMZ-20566); *S. pseudopneumoniae* (DSMZ-18670); *S. pyogenes* (DSMZ-20565); *S. sanguinis* (DSMZ-20567); and *S. mitis* (DSMZ-12643)). Analysis was performed in QIIME^27^, using a clustering level of 95% to define OTUs. We attributed the most common OTU sequences to Streptococcal species by BLAST searches. Full details are online (http://hdl.handle.net/10044/1/63937).

### Statistical analysis

We used the Differential Expression Analysis for Sequence Count Data (DESeq2 function in R)^29^ to compare OTU abundance between subject and control groups, controlling the false positive rate at *P =* 0.05. Parameters extracted for each OTU included log_2_(fold change), globally adjusted *P* value and abundance and prevalence information. Two-sided *P* values are reported throughout.

Co-abundance networks between non-rarefied OTU abundances were analyzed using the WGCNA package^30^. Abundances were log transformed with 0.1 added to zeroes^31^, and the topological adjacency matrix was constructed from Spearman’s correlation coefficients with a *β* soft thresholding parameter of 3. Hierarchical clustering of the overlap matrix with dynamic tree cutting defined the co-abundance modules, with a minimum module size set at 20 OTUs.

The significance of Spearman’s correlation between module eigengenes and clinical variables was adjusted for multiple testing using the Benjamini and Hochberg method^32^. Module structure was contrasted between cohorts using the R package circlize (0.4.5).

## Results

### Structure of the normal airway microbiome

We submitted oropharyngeal swabs from 578 subjects to 16S rRNA gene qPCR and sequencing, the latter yielding 44,290,100 high quality reads (Supplementary Figure 1 for analysis flowchart). After removal of 173 OTUs with high probability of being contaminants and 13,472 rare OTUs present in only one sample or with less than 20 reads, there remained 4,218 OTUs derived from 43,775,771 reads. To enable diversity analyses based on proportions, the samples were rarefied to a minimum of 6,543 reads, retaining 529 samples containing 4,005 OTUs and 3,461,247 reads. For consistency, unrarefied data from these same 529 samples were used to test differences between subject groups by DeSEQ2, and network analyses. No systematic differences in results were seen if the larger sample was analyzed.

The average age of the 529 subjects was 56 years (Supplementary Table 1). Sixty subjects were current smokers and 216 were ex-smokers (with a mean 18 years since quitting). The mean levels of the forced expiratory volume in one second (FEV1) and the forced vital capacity (FVC) of the subjects were normal. There were 77 doctor-diagnosed asthmatics, 82% of whom were atopic by prick skin tests (47% of the rest of the population were also atopic). There was only one case with a clinical diagnosis of COPD. The frequency of asthma and current smoking were not different to the whole Busselton cohort^16^.

An estimate of Bray Curtis beta diversity (β) for the population gave the mean dissimilarity in microbial diversity (M) between subjects to be 0.51 ± SD=0.06 (on a scale of 0-1), indicating that on average individual airway microbiomes shared half of their OTUs.

Five phyla contained 98.4% of all OTUs (Table 1, Supplementary Table 2). Firmicutes (predominately *Streptococcus* and *Veilonella* spp.) was the most common phylum, with 24 OTUs in the top 50, and 57.9% of all OTUs found in the complete dataset. Bacteroidetes (predominately *Prevotella* spp.) contained 14.1% of the OTUs, Proteobacteria (predominately *Neisseria* and *Haemophilus* spp.) contained 12.3%, Actinobacterium 9.1% and Fusobacterium 4.9%. Overall, the 50 most abundant OTUs accounted for 92% of the data (Supplementary Table 2).

**Table 1.**
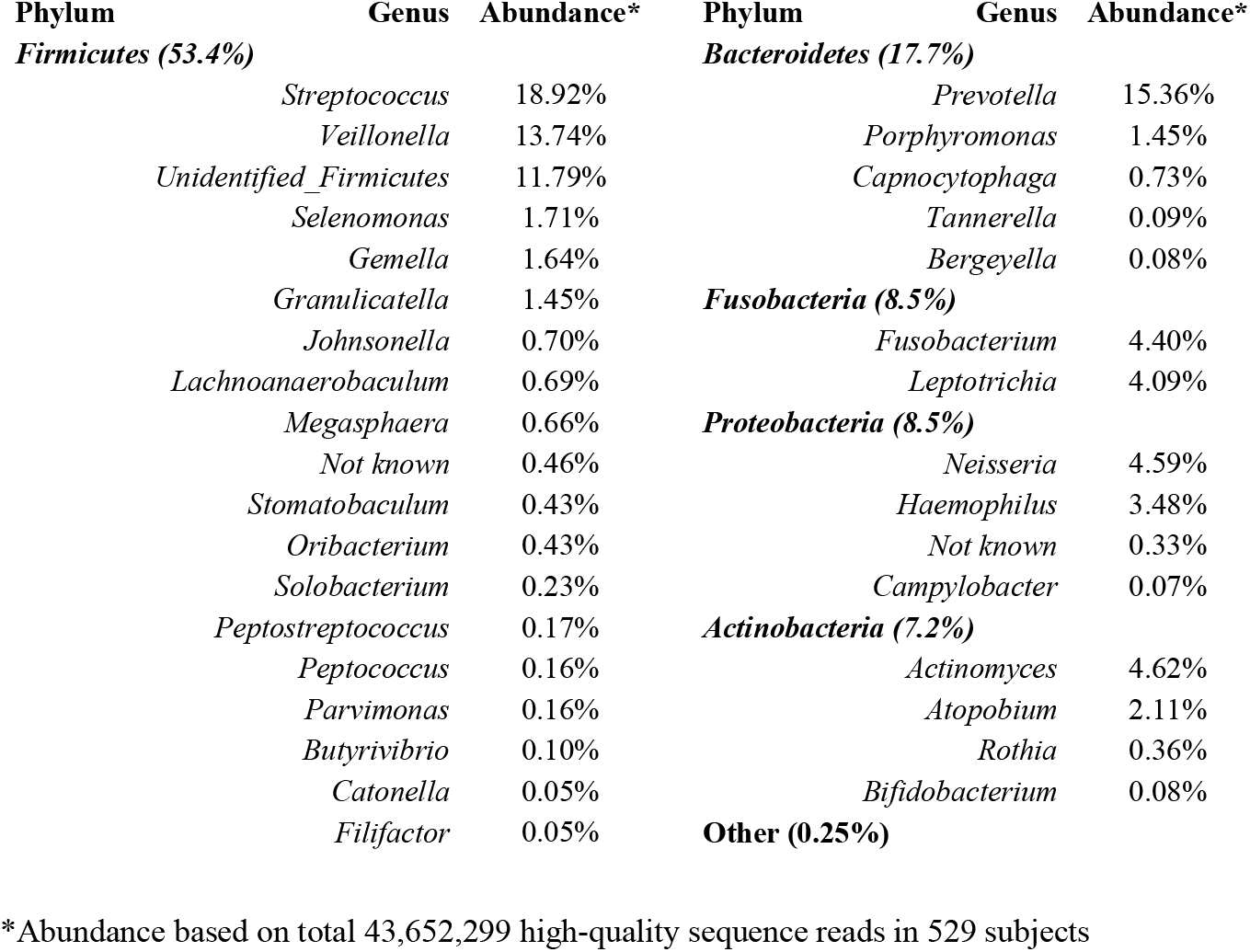
Principal phyla and genera of airway bacteria in a general population sample

We sequenced the *map* gene to differentiate between *Streptococcus* taxa^20^ in 483 subjects. After removal of sequences only present in one sample or with fewer than 20 reads or negative correlations with qPCR abundance there remained 14,898 map_OTUs (Supplementary Figure 2), indicating very substantial variation in Streptococcal strains in the population. β diversity estimates in rarefied data (to a level of 7,700 reads) found M = 0.84 ± SD=0.06, indicating low similarity of the streptococcal composition between subjects. The nine most prevalent *map*_OTUs were identified as *S. salivarius*, with *S. parasanguinis* the tenth most prevalent. (Supplementary Table 7). The potential pathogen S. *mitis/pneumoniae* was detected in 58% of subjects, although at low abundance.

Following network analyses with WGCNA^22^ we observed 13 discrete modules in which the abundance of members was strongly correlated. Just 13 OTUs remained unassigned to a network. The WGCNA program labels modules with unique colour identifiers, and we have named them according to their most abundant genera (Table 2). OTUs unassigned to a network are referred to as the grey module. The 5 largest modules (in terms of abundance of members) contained contained 97.6% of all OTU sequence reads (Table 2).

**Table 2.** WGCNA module summary and associations with smoking

| Module ID<br>(Colour) | Number<br>of OTUs | Total<br>Abundance | Overall<br>% | Cum<br>% | Smoking<br>R | Smoking<br>P | Module description |
| --- | --- | --- | --- | --- | --- | --- | --- |
| **Prevotella.1**<br>(Turquoise) | 2218 | 18,636,985 | 42.69 | 42.69 |  |  | Commensal carpet: *Veillonella*, *Prevotella*, *Actinomyces*. *Veillonella* and *Atopobium* hubs |
| **Streptococcus.2**<br>(Blue) | 472 | 9,433,313 | 21.61 | 64.30 | -0.13 | 2.E-02 | *Streptococcus* and *Haemophilus* prevalent. *Lactobacilliae* and *Gemella* hubs |
| **Streptococcus.1**<br>(Magenta) | 126 | 8,480,289 | 19.43 | 83.73 | 0.18 | 8.E-04 | *Streptococci* dominated |
| **Fusobacteria**<br>(Brown) | 583 | 3,099,110 | 7.10 | 90.83 | -0.26 | 4.E-08 | *Fusobacteria* and *Leptotrichia* hubs |
| **Neisseria**<br>(Green) | 204 | 2,969,651 | 6.80 | 97.63 | -0.35 | 1.E-14 | *Neisseria* dominated, prevalent *Capnocytophaga* |
| **Prevotella.2**<br>(Black) | 136 | 387,098 | 0.89 | 98.52 | 0.15 | 4.E-03 | *Prevotella*, *Parvimonas*, *Streptococci*, *Porphyromonas* |
| **Veillonella**<br>(Cyan) | 50 | 173,186 | 0.40 | 98.92 | 0.17 | 1.E-03 | *Veillonella* |
| **Prevotella.3**<br>(Purple) | 71 | 105,630 | 0.24 | 99.16 |  |  | *Prevotella* dominated |
| **Indeterminate**<br>(Tan) | 55 | 101,284 | 0.23 | 99.39 | 0.15 | 4.E-03 | *Prevotella* and *Treponema* |
| **Porphyromonas**<br>(Salmon) | 50 | 89,951 | 0.21 | 99.60 | 0.15 | 6.E-03 | *Porphyromonas* and *Prevotella* |
| **Bifidobacteria**<br>(Pink) | 134 | 86,562 | 0.20 | 99.80 | 0.32 | 8.E-13 | *Bifidobacterium* hubs |
| **Peptococcus**<br>(Midnightblue) | 44 | 79,920 | 0.18 | 99.98 | -0.16 | 2.E-03 | *Peptococcus* |
| **Contaminants**<br>(GreenYellow) | 62 | 8,869 | 0.02 | 100.00 | -0.12 | 3.E-02 | *Herbaspirillum*: potential contaminants |
| **Unconnected**<br>(Grey) | 13 | 451 | 0.00 |  |  |  | Unconnected OTUs: potential contaminants |

Individual hubs were very strongly connected to their network vectors (range of *P* = 7.9E-266 to 1.9E-121) (Supplementary Table 4), and the strengths of association suggest that these coabundance modules represent “guilds” of co-operating bacteria that may occupy ecological niches on the mucosa.

The largest guild *(Prevotella.1:* turquoise) accounted for 42.7% of reads (Table 2, Supplementary Table 4). The most common organisms were within the genera *Prevotella, Veillonella, Actinomyces* and *Atopobium.* These organisms resemble common mucosal commensals at other body sites, and perhaps represent a base microbial carpet. The smaller guild (cyan) on the same division (B) of the network dendrogram (Supplementary Figure 3) was almost entirely made up of *Veillonella* spp. and may occupy a related ecological niche.

The *Streptococcus.2* module (blue) contained 21.6% of reads, predominately from the genera *Streptococcus, Haemophilus* and *Veillonella.* Network hubs included *Lactobacillales* and *Gemella.* The adjacent network *(Neisseria:* green) (Supplementary Figure 3) was dominated by *Neisseria*, with *Porphyromonas*, and *Capnocytophagia*. This may suggest a normal guild that can be occupied by *Proteobacteria* potential pathogens.

The *Streptococcus.1* module (magenta) (19.4% of reads) was completely dominated by *Streptococcus* taxa (40%) and an unidentified *Firmicutes* (60%) (Supplementary Table 4) which is likely also to be streptococcal (based on phylogenetic clustering, not shown). Network hubs were also *Streptococcus*.

### Smoking

A stepwise regression (IBM SPSS Statistics v25) found that microbial diversity in individual airways was independently related to current cigarette smoking (R^2^=6%, P<0.001), a current diagnosis of asthma (additional R^2^=1.4%, P<0.005) and packyears of smoking (additional R^2^=0.8%, P=0.04) (Supplementary Table 3), but not to age or sex. We therefore partitioned the data into three subgroups: smoking + packyears>10 (n=159); asthmatic (n=77); and unaffected (n=233).

A DeSEQ2 analysis to identify significant differences in the abundance of specific taxa revealed marked effects of cigarette smoking. (Figure 1, Supplementary Figure 4, Supplementary Table 5a and 5b). The loss of diversity affected many abundant OTUs, including those in the genera *Fusobacterium*, *Neisseria*, *Haemophilus*, *Veillonella* and *Gemella.* By contrast, the OTUs increased in smokers were in general highly abundant *Streptococci.* Examination of *map* gene OTUs attributed increases in abundance to *S. parasanguinis* (log_2_(Fold change) 5.2, *P_adjusted_=1.75E-07), S. mitis/pneumoniae* (3.62, 4.81E-09), *S. salivarius* (3.03, 5.59E-15) and *S. thermophilus* (2.53, 7.38E-05) (Supplementary Table 8).

**Figure 1.**
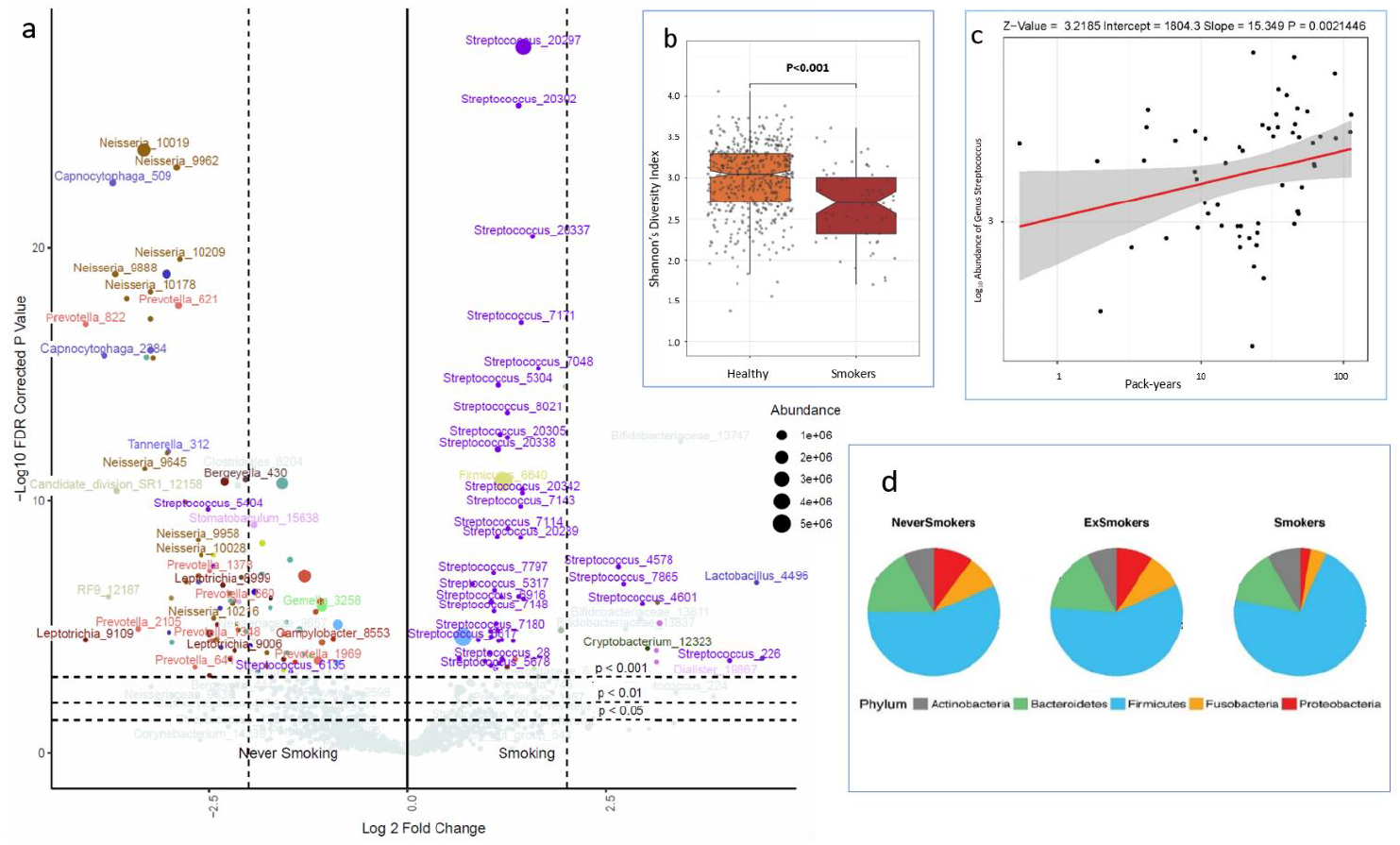
Smoking and the airway microbiome. **a)** The volcano plot shows significant differences in the abundance of OTUs between current smokers and the rest of the population. Fold change is shown on the x axis and −log10 *P* (FDR corrected) on the y axis. Relative abundances are reflected in the data point sizes; **b)** shows differences in alpha diversity between smokers and never smokers (boxes show inter-quartile range, notches 95% CI of the median, *P* values are two-sided from multiple regression); **c)** shows progressive increase in the abundance of *Streptococcus* OTUs with increasing packyears (shaded area indicates 95% CI); and **d)** shows how the phyla of ex-smokers resemble never-smokers, implying a beneficial effect of smoking cessation.

To further explore the impact of smoking and asthma on the higher order structure of the airway microbiome, co-abundance networks were constructed separately in the asthmatic and current smoker portions of the cohort and compared with the full dataset (representing the whole population) (Supplementary Figure 4). The analysis was limited to the 4,207 OTUs present in all three datasets.

The network structure of the communities was profoundly altered in current smokers. Whilst the largest guild (Prevotellla.1: commensal carpet) showed relative preservation, other modules showed markedly lower levels of conservation and were strongly positively or negatively associated with smoking status; either in terms of module eigenvectors or hubs (Figure 2, Table 2, Supplementary Table 4). In smokers, 276 OTUs became disconnected from any module. These most strongly featured *Streptococcus* (70 OTUs), unknown genera (41 OTUs) and *Veillonella* (35 OTUs).

**Figure 2.**
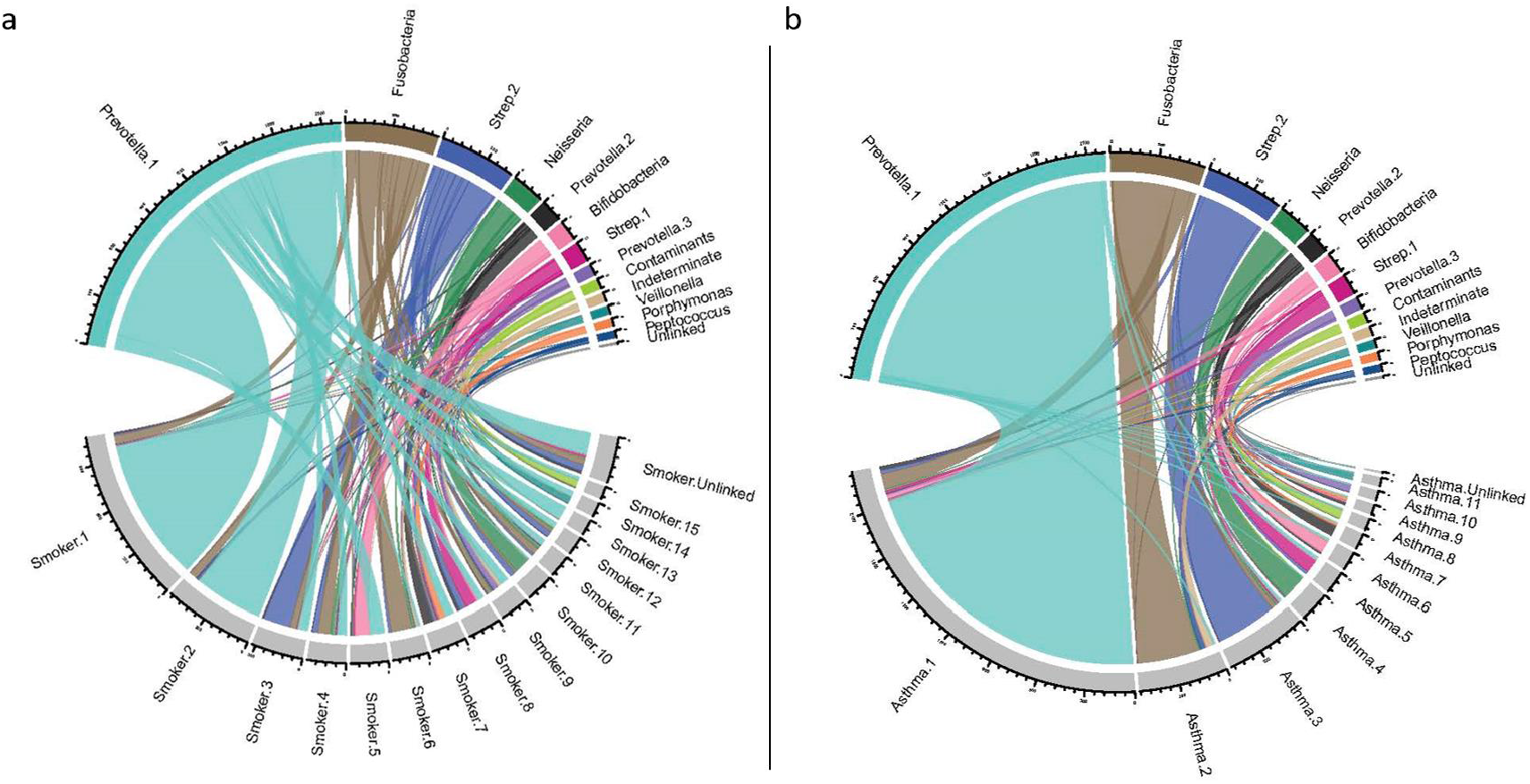
Network structure of the airway microbiome in normal subjects, compared to smokers and asthmatics. The Chord plots show sharing and discordance of 4,207 OTUs common to the three datasets for co-abundance networks. **a)** Network membership in the whole population (top half of plot) compared to current smokers (bottom half of plot); and **b)** compared to asthmatics. Module colours are arbitrarily assigned by WGCNA, and module bacterial names are derived from Table 2. Modules in smokers and asthmatics are simply named by size (Smoking.1, Asthma.2, etc.). There is a marked change of structure with fragmentation of major networks in the smokers, but high conservation of network membership between asthmatics and the whole cohort.

### Asthma

Microbial diversity loss in asthmatics compared to non-smoking subjects was qualitatively different to the effects of smoking. DeSEQ analysis showed only two taxa *(Neisseria* and *Rothia* OTUs) to be increased in abundance in asthmatic airways *(P_adjusted_<0.05)* (Figure 3, Supplementary Table 6a). Of these, the *Neisseria* OTU was abundant (4.7% of reads in the population) and showed a 2-fold increase, consistent with increases in *Protebacteria* spp. consistently observed in excess by comparisons of asthmatic and normal airways^2,3,33,34^.

**Figure 3.**
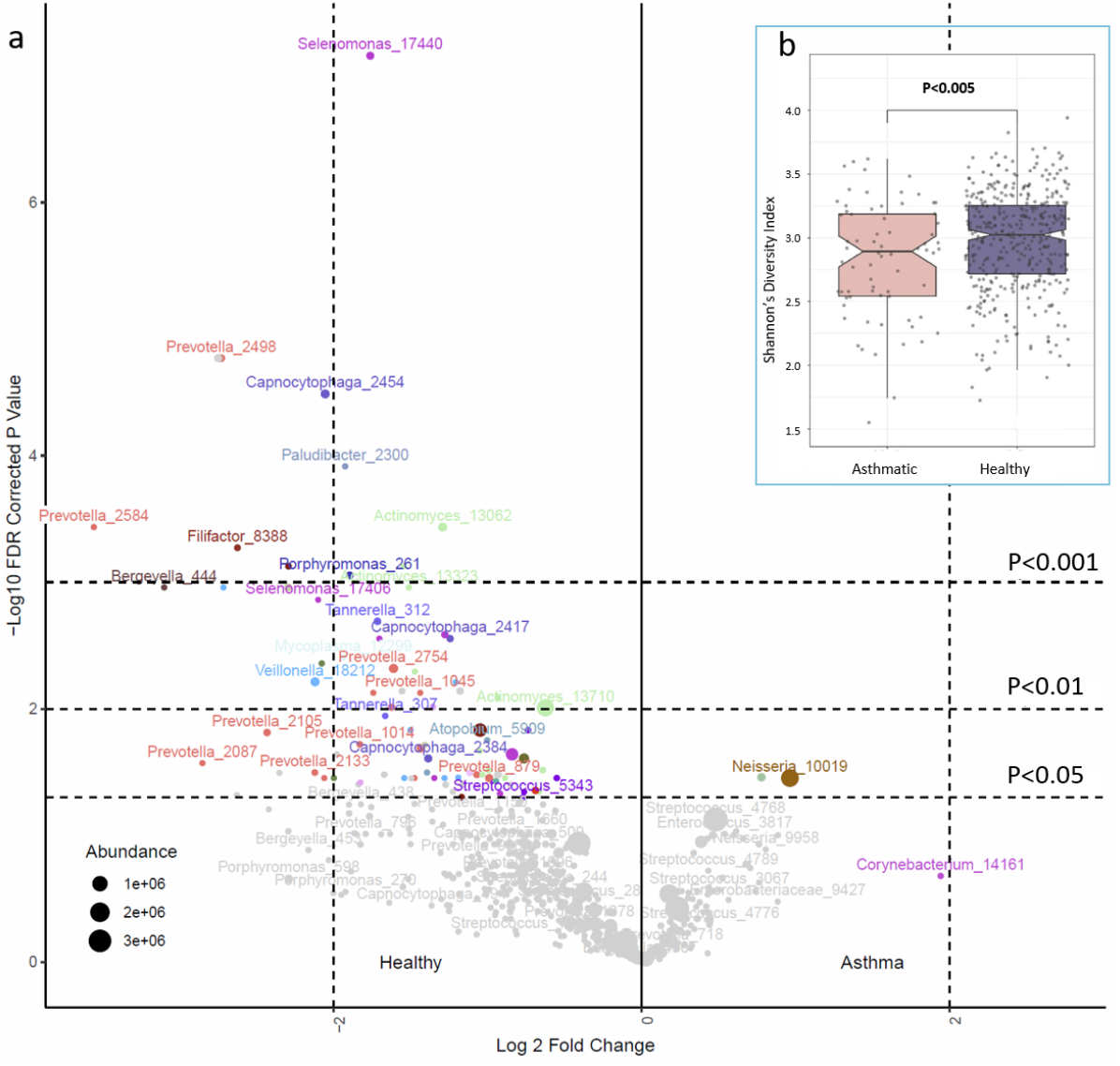
Asthma and the airway microbiome. **a)** The volcano plot shows significant differences in the abundance of OTUs between asthmatics and non-smoking subjects with less than 10 packyears of lifetime exposure. Fold change is shown on the x axis and −log10 *P* (FDR corrected) on the y axis. Relative abundances are reflected in the data point sizes; **b)** shows differences in alpha diversity between asthmatics and unaffected non-smoking subjects (boxes show inter-quartile range, notches 95% CI of the median, *P* values are two-sided from multiple regression).

Eighty-four OTUs were in relatively low abundance amongst asthmatic subjects (Figure 3, Supplementary Table 6b). In marked contrast to smokers, the affected organisms were often in poorly characterized or potentially fastidious genera, including *Leptotrichia*, *Selenomonas*, *Megasphaera* and *Capnocytophaga*. Some representatives of the more common genera *Actinomyces, Prevotella* and *Veillonella* were also less abundant.

Inhaled corticosteroids (ICS) are widely used in the maintenance treatment of asthma, and 51 (66%) of our asthmatics were currently using such therapy. Logistic regression analyses showed no independent effect on ICS use from OTUs positively or negatively associated with asthma, or with microbial diversity.

The module eigenvectors did not correlate with the presence of asthma, indicating that the general structure of oropharyngeal microbial communities in asthmatics was preserved (Figure 2). Nevertheless, the asthma-enriched *Neisseria_10019* taxon was a hub of the Neisseria guild, which also contained the significantly reduced *Capnocytophagia_2454* (Supplementary Table 4). Other asthma-reduced taxa were concentrated in the Prevotella.1 (containing 57 of the 84 asthma-associated OTUs) and Prevotella.2 (12/84) guilds (Chi^2^ exact test, P=2.8×10^−8^). Asthma-associated OTUs were enriched amongst the most highly connected module members (OR=18.6, P=2.9×10^−9^), and so are well positioned to influence host-microbial interactions. The Neisseria, Prevotella.1 and Prevotella.2 guilds thus provide a focus for further understanding of the ecology of asthmatic airway microbiota.

## Discussion

Our study shows that the healthy airway microbiota are contained within a structured ecosystem that is conserved across the general population. In this aspect are they are thus similar to commensals at other body sites. The main phyla in airway samples (Firmicutes, Bacteriodetes, Actinobacteria, and Proteobacteria) also dominate the gut^35^, skin^36^ and vagina^37^, although bacterial genera and species differ considerably between body surfaces, Our results were well powered to map microbial community composition, but we could only surmise limited functions by genus assignments and network relationships. Metagenomic shotgun sequencing has given profound insights into the function of the gut microbiota, but shotgun sequencing has been problematic for respiratory samples because high concentrations of human DNA (~99%) interfere with PCR and require a 100 fold increase in sequencing depth (and cost) to derive coverage comparable to faecal samples^4^. Additionally, genome assembly from shotgun sequences is limited by a paucity of reference genomes for airway commensals. Secreted host factors that either constrain airway pathogens or support commensal bacteria are known to exist^38^, but they have also not yet been systematically surveyed.

Cigarette smoking is known to affect the airway microbiota^7^, but the extent and specificity of disruption shown here suggests an independent capacity to damage human health. The loss of diversity may predispose smokers to the recurrent infections that lead to COPD^6,7^. The annual rate of antibiotic prescription in the Australian population is 254 per 1000, and half of these will be for respiratory infections^39^, so it is likely that many smokers will have intermittently been given antibiotics which will have contributed to the microbial community abnormalities. Smoking is accompanied by substantial changes in the bowel flora^40^ that may mediate smoking influences on inflammatory bowel disease. Bacteria have known roles in the genesis of cancer in general^41^ and in lung cancer specifically^8^. *Streptococcus* spp. produce an array of potent toxins that act against human cells or tissues^42^, and the expansion of *Streptococcus* clades in smokers might be carcinogenic. Most patients with lung cancer have been heavy smokers and smoking often continues after diagnosis. Our results might also suggest that the local lung microbiota should be considered a factor in lung cancer responses to immunotherapy^43^.

Although the profound consequences of cigarette smoking are clear, the community degradation seen in asthmatics is more subtle and without an obvious cause. Importantly, we have been able to show that ICS are not associated with an abnormal microbiome, consistent with published results^44^. Asthma is not considered an indication for antibiotics in the Australian healthcare system. A simple iatrogenic pruning of diversity in asthmatics therefore appears improbable.

Divergent (but potentially complementary) theories are offered on mechanisms by which microbial diversity might prevent asthma. The “immune deviation” hypothesis suggests that the adaptive immune system needs exposure to infections in order to avoid inappropriate reactions^45^. An extension of this model is that absence of commensal organisms leads to loss of local or systematic tonic signals that normally down-regulate immune responses at mucosal surfaces^46^. Our findings, of reduced numbers of distinctive low-abundance organisms, are consistent with immune modulation by these organisms.

However, the consistent finding of excesses of *Proteobacteria* in this and other studies ^2,3,34^ (and *Streptococcus* spp. in severe disease ^2,47,48^) are also consistent with asthmatic airway inflammation that follows intermittent mucosal damage by bacteria. *Proteobacteria* include many known potential pathogens from the genera *Haemophilus*, *Moraxella*, and *Neisseria* that, despite the ability to cause disease, are commonly carried without symptoms in the population (“pathobionts”)^4^. In the “asthma as an infection” hypothesis it becomes possible that a diverse microbial community protects against asthma through inhibition of pathobiont effects, by modifying their growth, adherence or biofilm formation^49^.

Our results provide a strong impetus to isolate and study the organisms that are perturbed in asthmatic airways, and to test hypotheses that involve immune modulation or mucosal damage.

The demonstration of a highly ordered and conserved microbiome is relevant to many lung disorders^4^, particularly the respiratory infections that take 4 million lives annually^50^. Factors that trigger the switch between colonisation and infection include the density of pathobionts in vulnerable sites, synergism and nutrient competition with commensals^51^, and intercurrent viral infections. Diversity in the gut microbiota confers colonization resistance to intestinal infections^52–54^, and our results allow consideration of therapeutic manipulation of the pulmonary microbiome.

## Supporting information

Turek supplementary information

## Acknowledgements

The study was funded by the Asmarley Trust and a Wellcome Senior Investigator Award to WOCC and MFM (P46009). The Busselton Healthy Ageing Study is supported by grants from the Government of Western Australia (Office of Science, Department of Health) and the City of Busselton, and from private donations to the Busselton Population Medical Research Institute. We thank the WA Country Health Service and the community of Busselton for their ongoing support and participation.

## Author contributions

Overall study design: AWM, AJ, MH, JH, MJC, MFM, and WOCMC. Busselton Survey and sample collection: MH, JH, AJ, AWM. Microbial analysis strategy (laboratory and bioinformatics); MJC, MFM, PJ, EMT. Laboratory experiments: EMT, with assistance and advice by MJC, PJ, LC and MFM. Primary ecological analyses EMT with input from MJC and PJ. Network analysis EMT and SWO. Secondary analyses WOCC. EMT, MFM and WOCMC wrote the first draft of the paper. All authors have read and contributed to the final version of the paper.

## Materials & Correspondence

The raw data is available online at the European Nucleotide Archive at the European Bioinformatics Institute, with the accession number PRJEB29091

The R scripts for analysis are available at https://tinyurl.com/y2onjblt

