## Supplementary material for "Smoking, asthma and airway microbial disruption": Turek supplementary information

### Supplementary Information Turek *et al.* 2020

#### Contents

Figures ... 2

Supplementary Figure 1. Flow diagram showing stages of data clean-up and analyses for 16S rRNA gene sequences ... 2

Supplementary Figure 2. Flow diagram showing stages of data clean-up and analyses for *map* sequences to distinguish *Streptococcus* spp. ... 2

Supplementary Figure 3. Composition diagram showing subjects for subgroup analyses of smoking and asthma associations. ... 4

Tables ... 5

Supplementary Table 1. Subject characteristics for the 529 individuals with completed 16S analyses ... 5

Supplementary Table 2. Top 100 OTUs in all subjects ... 6

Supplementary Table 3. Stepwise regression for determinants of Shannon alpha diversity (IBM SPSS 25) ... 10

Supplementary Table 4. Weighted Gene Correlation Network Analysis (WGCNA): Principal Module Membership and Hubs ... 11

Supplementary Table 5a. Individual OTUs increased in smokers ... 20

Supplementary Table 5b. OTUs decreased in smokers ... 22

Supplementary Table 6a. OTUs increased in asthmatics ... 25

Supplementary Table 6b. OTUs decreased in asthmatics ... 26

Supplementary Table 7. Analysis of *map* gene: frequencies and identities of *Streptococcus* spp. ... 29

Supplementary Table 8. *Streptococcus* spp. affected by smoking ... 31

### Figures

#### Supplementary Figure 1. Flow diagram showing stages of data clean-up and analyses for 16S rRNA gene sequences

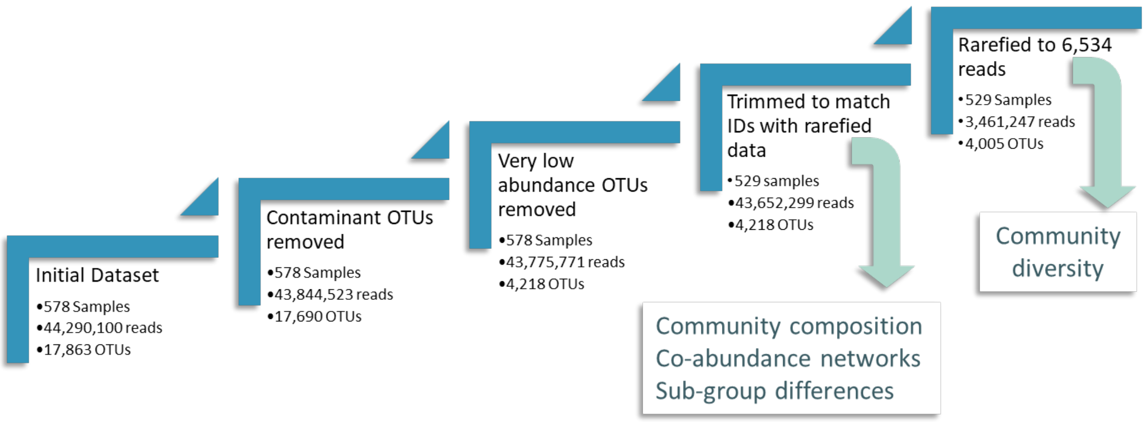

The diagram reads upward from left to right. OTU=operational taxonomic unit, based on 16S RNA gene sequences.

#### Supplementary Figure 2. Flow diagram showing stages of data clean-up and analyses for *map* sequences to distinguish *Streptococcus* spp.

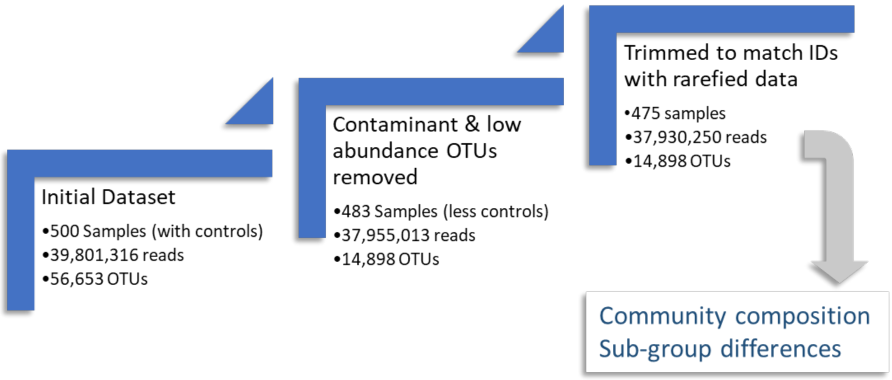

The diagram reads upward from left to right. OTU=operational taxonomic unit, based on *map* gene sequences.

### Supplementary Figure 3. Characteristics of the airway microbiome in a general population

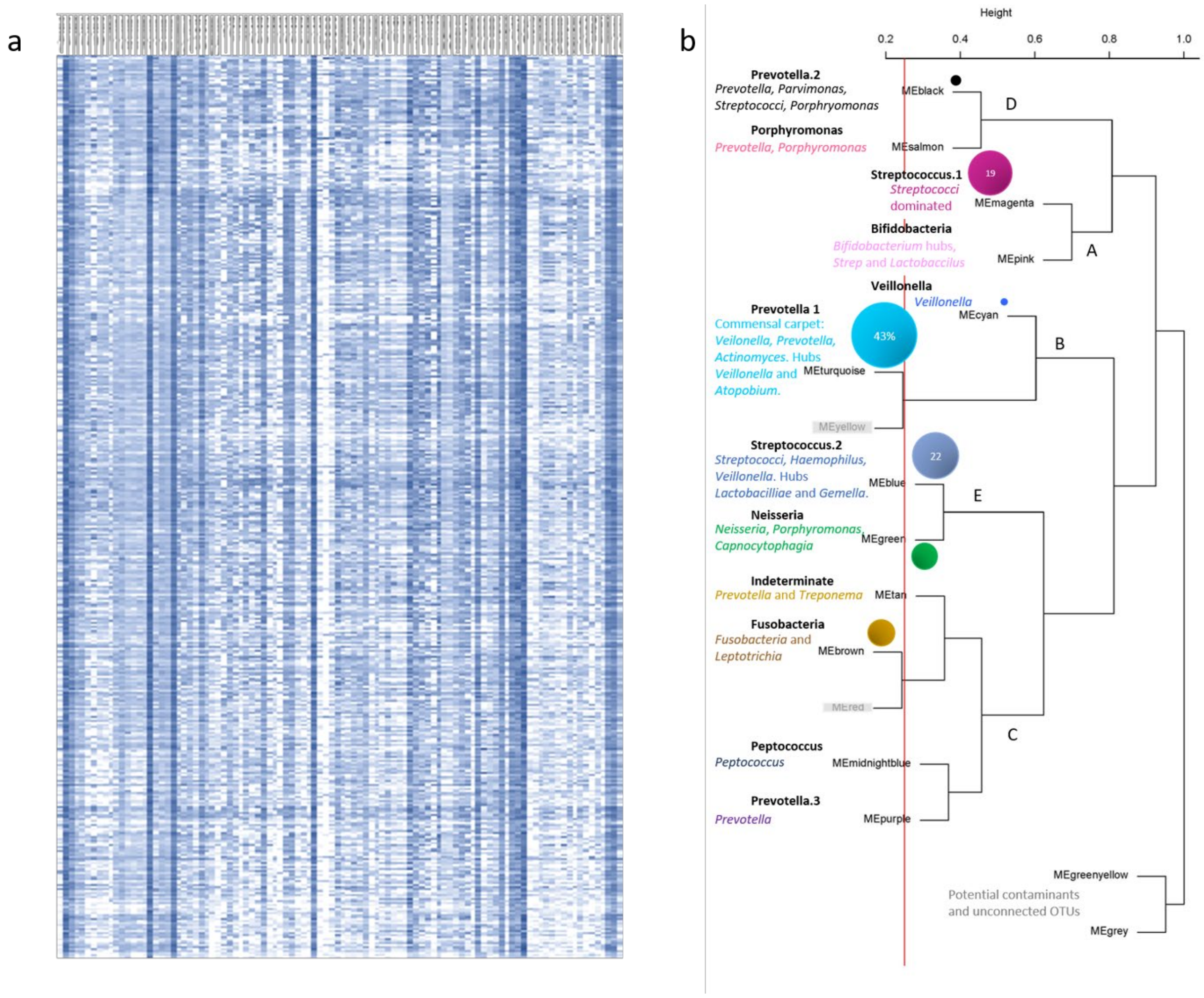

a) Heatmap of the log abundance of the top 100 taxa (OTUs from 16S rRNA sequences) in all subjects. OTUs are shown on the x axis and individual subject results along the y. Community composition is conserved across the population; b) Relationship between WGCNA networks, based on correlation between neighbouring module members. Red and Yellow modules are below the differentiation threshold (red line) and are merged with their immediate neighbours (turquoise and brown respectively). Bacteria not connected to other taxa are in the grey module. Members of the greenyellow module include known contaminants, consistent with the distance from other networks;

Supplementary Figure 4. Composition diagram showing subjects for subgroup analyses of smoking and asthma associations.

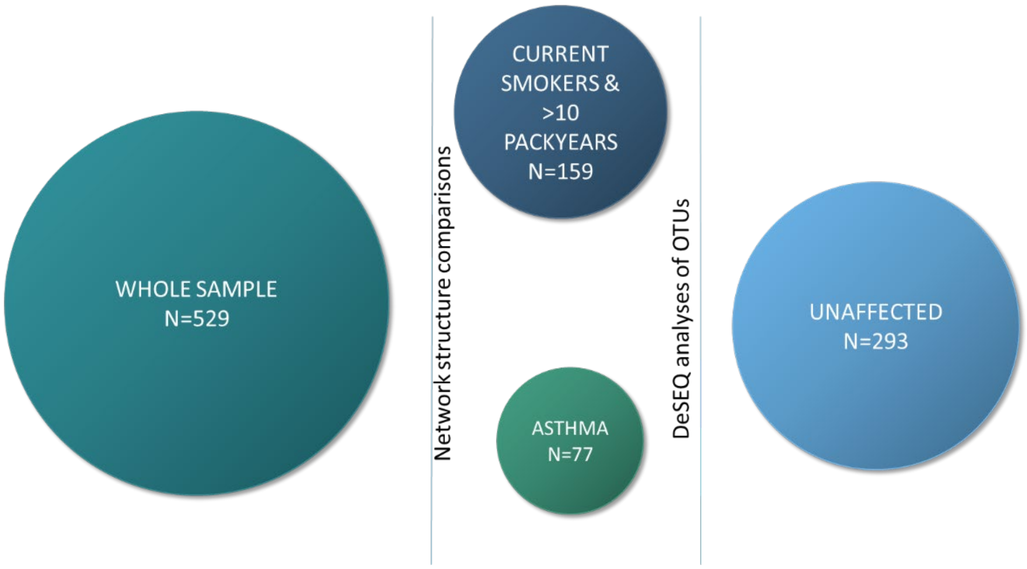

The diagram shows the subjects used in analysing associations with smoking and asthma. WGCNA networks were generated on the whole sample (representing the Busselton population), and then compared with networks from heavy smokers and from asthmatics. To examine OTU associations with disease, abundance differences were tested between smokers or asthmatics against subjects unaffected by these conditions.

### Tables

#### Supplementary Table 1. Subject characteristics for the 529 individuals with completed 16S analyses

| | Male | Female |
|---|---|---|
| Total | 259 | 270 |
| Age **(years)** | 55.71 (SD:5.57) | 55.78 (SD:5.59) |
| BMI | 28.48 (SD:4.04) | 27.28 (SD:5.07) |
| Current Smoker | 35 | 25 |
| Number of Pack Years | 11.83 (SD:18.70) | 6.53 (SD:12.79) |
| Ex-Smoker | 108 | 108 |
| Years since Quit | 17.52 (SD:12.76) | 20.21 (SD:11.31) |
| Never Smoker | 116 | 137 |
| Asthma | 36 | 41 |
| Atopy (non-asthma) | 55% | 39% |
| Atopy (asthmatics) | 88% | 76% |
| ppFEV baseline | 96.41 (SD:13.98) | 94.79 (SD:12.67) |
| ppFVC baseline | 98.92 (SD:11.95) | 98.55 (SD:12.04) |

BMI = Body Mass Index, Atopy = 3mm or greater skin prick test response to mixed grasses or to *D. pteronyssinus* or *D. farinae*, ppFEV baseline = percent predicted forced expiratory volume baseline, ppFVC baseline = percent predicted forced vital capacity baseline

#### Supplementary Table 2. Top 100 OTUs in all subjects

| Row Labels | Total Sequences (Abundance) | Prevalence (out of 529 subjects) |
|---|---|---|
| ***Firmicutes*** | **23373367** | |
| ***Streptococcus*** | **8258368** | |
| *Streptococcus_20297* | 3019924 | 529 |
| *Streptococcus_20302* | 39744 | 528 |
| *Streptococcus_20338* | 61991 | 527 |
| *Streptococcus_28* | 62335 | 520 |
| *Streptococcus_4768* | 4896262 | 529 |
| *Streptococcus_6617* | 14951 | 517 |
| *Streptococcus_6789* | 23088 | 498 |
| *Streptococcus_6815* | 18701 | 500 |
| *Streptococcus_6916* | 22255 | 525 |
| *Streptococcus_7798* | 64585 | 529 |
| *Streptococcus_7928* | 34532 | 514 |
| ***Veillonella*** | **5999024** | |
| *Veillonella_16405* | 23284 | 451 |
| *Veillonella_16908* | 19869 | 470 |
| *Veillonella_18212* | 163662 | 380 |
| *Veillonella_19164* | 23913 | 455 |
| *Veillonella_19388* | 97522 | 526 |
| *Veillonella_19389* | 89839 | 527 |
| *Veillonella_19390* | 817686 | 529 |
| *Veillonella_19412* | 4763249 | 529 |
| ***Unidentified_Firmicutes*** | **5146895** | |
| *Firmicutes_6640* | 5146895 | 529 |
| ***Selenomonas*** | **745942** | |
| *Selenomonas_17440* | 67889 | 525 |
| *Selenomonas_17559* | 643883 | 529 |
| *Selenomonas_17724* | 34170 | 486 |
| ***Gemella*** | **714904** | |
| *Gemella_3258* | 714904 | 529 |
| ***Granulicatella*** | **633823** | |
| *Granulicatella_3873* | 24379 | 441 |
| *Granulicatella_3979* | 609444 | 529 |
| ***Johnsonella*** | **306698** | |
| *Johnsonella_15517* | 306698 | 495 |
| ***Lachnoanaerobaculum*** | **299622** | |
| *Lachnoanaerobaculum_14874* | 224079 | 527 |
| *Lachnoanaerobaculum_14972* | 75543 | 516 |
| ***Megasphaera*** | **287130** | |
| *Megasphaera_16215* | 287130 | 528 |
| ***NA*** | **202588** | |
| *Clostridiales_8242* | 16636 | 394 |

| | | |
|---|---|---|
| *Lactobacillales_3640* | 14800 | 502 |
| *Lactobacillales_3650* | 21695 | 507 |
| *Ruminococcaceae_11741* | 23711 | 411 |
| *Ruminococcaceae_11808* | 35320 | 464 |
| *Veillonellaceae_17891* | 90426 | 410 |
| ***Stomatobaculum*** | **189635** | |
| *Stomatobaculum_15638* | 81904 | 481 |
| *Stomatobaculum_15766* | 107731 | 501 |
| ***Oribacterium*** | **185734** | |
| *Oribacterium_15113* | 40336 | 506 |
| *Oribacterium_15121* | 20558 | 391 |
| *Oribacterium_15965* | 124840 | 524 |
| ***Solobacterium*** | **102474** | **525** |
| *Solobacterium_11985* | 102474 | 525 |
| ***Peptostreptococcus*** | **74697** | |
| *Peptostreptococcus_20146* | 74697 | 486 |
| ***Peptococcus*** | **70217** | |
| *Peptococcus_9344* | 70217 | 466 |
| ***Parvimonas*** | **68222** | |
| *Parvimonas_8503* | 68222 | 513 |
| ***Butyrivibrio*** | **45790** | |
| *Butyrivibrio_14629* | 45790 | 438 |
| ***Catonella*** | **21066** | |
| *Catonella_9245* | 21066 | 495 |
| ***Filifactor*** | **20538** | |
| *Filifactor_8388* | 20538 | 299 |
| ***Bacteroidetes*** | **7726297** | |
| ***Prevotella*** | **6705316** | |
| *Prevotella_1014* | 18947 | 265 |
| *Prevotella_1525* | 4610220 | 529 |
| *Prevotella_1736* | 28316 | 434 |
| *Prevotella_1750* | 16281 | 397 |
| *Prevotella_1791* | 25413 | 462 |
| *Prevotella_1868* | 351150 | 526 |
| *Prevotella_1969* | 346641 | 519 |
| *Prevotella_2105* | 33863 | 214 |
| *Prevotella_2177* | 22805 | 456 |
| *Prevotella_2498* | 45434 | 380 |
| *Prevotella_2598* | 685534 | 527 |
| *Prevotella_2754* | 168126 | 476 |
| *Prevotella_2890* | 89671 | 446 |
| *Prevotella_621* | 144656 | 495 |
| *Prevotella_822* | 29280 | 376 |
| *Prevotella_879* | 88979 | 494 |
| ***Porphyromonas*** | **631475** | |
| *Porphyromonas_11896* | 491972 | 519 |

| | | |
|---|---|---|
| *Porphyromonas_261* | 54868 | 420 |
| *Porphyromonas_598* | 84635 | 87 |
| ***Capnocytophaga*** | **317872** | |
| *Capnocytophaga_2384* | 64767 | 399 |
| *Capnocytophaga_2417* | 26121 | 437 |
| *Capnocytophaga_2454* | 113957 | 483 |
| *Capnocytophaga_509* | 113027 | 495 |
| ***Tannerella*** | **38759** | |
| *Tannerella_312* | 38759 | 430 |
| ***Bergeyella*** | **32875** | |
| *Bergeyella_430* | 32875 | 490 |
| ***Fusobacteria*** | **3706367** | **2417** |
| ***Fusobacterium*** | **1921431** | |
| *Fusobacterium_7409* | 1921431 | 529 |
| ***Leptotrichia*** | **1784936** | |
| *Leptotrichia_8776* | 866757 | 527 |
| *Leptotrichia_8876* | 215430 | 362 |
| *Leptotrichia_8929* | 377771 | 492 |
| *Leptotrichia_8995* | 324978 | 507 |
| ***Proteobacteria*** | **3692787** | |
| ***Neisseria*** | **2002143** | |
| *Neisseria_10019* | 1735113 | 525 |
| *Neisseria_10020* | 16590 | 397 |
| *Neisseria_10178* | 15187 | 420 |
| *Neisseria_9888* | 76462 | 428 |
| *Neisseria_9962* | 158791 | 502 |
| ***Haemophilus*** | **1517994** | |
| *Haemophilus_10797* | 96360 | 516 |
| *Haemophilus_10843* | 35829 | 507 |
| *Haemophilus_11056* | 165096 | 427 |
| *Haemophilus_11091* | 1199062 | 529 |
| *Haemophilus_11369* | 21647 | 392 |
| ***NA*** | **142834** | |
| *Neisseriaceae_10099* | 19372 | 434 |
| *Pasteurellaceae_11461* | 123462 | 515 |
| ***Campylobacter*** | **29816** | |
| *Campylobacter_8553* | 29816 | 517 |
| ***Actinobacteria*** | **3130379** | |
| ***Actinomyces*** | **2014748** | |
| *Actinomyces_13062* | 152157 | 509 |
| *Actinomyces_13710* | 1862591 | 529 |
| ***Atopobium*** | **921802** | |
| *Atopobium_5828* | 905892 | 529 |
| *Atopobium_5846* | 15910 | 454 |
| ***Rothia*** | **157077** | |
| *Rothia_13982* | 87838 | 524 |

| | | |
|---|---|---|
| *Rothia_14025* | 69239 | 406 |
| ***Bifidobacterium*** | **36752** | |
| *Bifidobacterium_13861* | 36752 | 249 |
| ***Candidate_division_TM7*** | **60470** | |
| ***NA*** | **60470** | |
| *Candidate_division_TM7_12053* | 60470 | 477 |
| ***Candidate_division_SR1*** | **49511** | |
| ***(blank)*** | **49511** | |
| *Candidate_division_SR1_12158* | 49511 | 341 |
| **Grand Total** | **41739178** | |

Supplementary Table 3. Stepwise regression for determinants of Shannon alpha diversity (IBM SPSS 25)

**Model Summary**

| | Change Statistics | | | | |
|---|---|---|---|---|---|
| Model | R Square Change | F Change | df1 | df2 | Sig. F Change |
| 1 | .060[a] | 33.005 | 1 | 515 | .000 |
| 2 | .014[b] | 7.905 | 1 | 514 | .005 |
| 3 | .008[c] | 4.323 | 1 | 513 | .038 |

a. Predictors: (Constant), Current Smoking

b. Predictors: (Constant), Current Smoking, Asthma

c. Predictors: (Constant), Current Smoking, Asthma, Packyears

**Coefficients[a]**

| Model | | Unstandardized Coefficients | | Standardized Coefficients | | | 95% Confidence Interval for B | |
|---|---|---|---|---|---|---|---|---|
| | | B | Std. Error | Beta | t | Sig. | Lower Bound | Upper Bound |
| 1 | (Constant) | 2.710 | .017 | | 155.278 | .000 | 2.675 | 2.744 |
| | Current Smoking | -.294 | .051 | -.245 | -5.745 | .000 | -.395 | -.194 |
| 2 | (Constant) | 2.729 | .019 | | 146.186 | .000 | 2.693 | 2.766 |
| | Current Smoking | -.301 | .051 | -.251 | -5.903 | .000 | -.401 | -.201 |
| | Asthma | -.131 | .047 | -.119 | -2.812 | .005 | -.223 | -.039 |
| 3 | (Constant) | 2.744 | .020 | | 137.569 | .000 | 2.705 | 2.783 |
| | Current Smoking | -.239 | .059 | -.199 | -4.071 | .000 | -.355 | -.124 |
| | Asthma | -.130 | .046 | -.119 | -2.809 | .005 | -.222 | -.039 |
| | Packyears | -.002 | .001 | -.102 | -2.079 | .038 | -.005 | .000 |

a. Dependent Variable: Shannon alpha diversity index

Supplementary Table 4. Weighted Gene Correlation Network Analysis (WGCNA): Principal Module Membership and Hubs

Colour shading for Abundance % (the % of reads for the OTU) is derived from the whole dataset and varies from green (highest) to red (lowest). Prevalence % is shown in blue (high) to red (low). Network hubs are highlighted in pink.

**TurquoiseModule: Prevotella.1**

| **OTU_Name** | **Genus** | **Abundance%** | **Prevalence%** | **-log10*(P)* MM** | **-log10*(P)* Smo** | **-log10*(P)* Asth** |
|---|---|---|---|---|---|---|
| Veillonella_19412 | Veillonella | 10.91 | 100.00 | 65.05 | | |
| Prevotella_1525 | Prevotella | 10.56 | 100.00 | 116.33 | | |
| Actinomyces_13710 | Actinomyces | 4.27 | 100.00 | 98.11 | | 2.01 |
| Atopobium_5828 | Atopobium | 2.08 | 100.00 | 111.68 | | |
| Leptotrichia_8776 | Leptotrichia | 1.99 | 99.62 | 83.12 | | 1.83 |
| Prevotella_2598 | Prevotella | 1.57 | 99.62 | 47.46 | | |
| Selenomonas_17559 | Selenomonas | 1.48 | 100.00 | 105.02 | | 1.64 |
| Leptotrichia_8929 | Leptotrichia | 0.87 | 93.01 | 48.22 | | |
| Prevotella_1868 | Prevotella | 0.80 | 99.43 | 130.59 | | |
| Prevotella_1969 | Prevotella | 0.79 | 98.11 | 73.41 | | |
| Megasphaera_16215 | Megasphaera | 0.66 | 99.81 | 115.22 | | 1.61 |
| Lachnoanaerobaculum_14874 | Lachnoanaerobaculum | 0.51 | 99.62 | 107.51 | | |
| Veillonella_19417 | Veillonella | 0.02 | 79.96 | 173.84 | | |
| Atopobium_5926 | Atopobium | 0.03 | 76.37 | 170.20 | | |
| Atopobium_5846 | Atopobium | 0.04 | 85.82 | 165.26 | | |
| Actinomyces_13214 | Actinomyces | 0.01 | 81.29 | 160.11 | | |
| Veillonella_19164 | Veillonella | 0.05 | 86.01 | 158.08 | | |
| Veillonella_16405 | Veillonella | 0.05 | 85.26 | 157.79 | | |
| Veillonella_16908 | Veillonella | 0.05 | 88.85 | 157.02 | | |
| Veillonella_18660 | Veillonella | 0.01 | 78.45 | 151.70 | | |
| Veillonella_16572 | Veillonella | 0.03 | 85.26 | 150.10 | | |
| Veillonella_17001 | Veillonella | 0.02 | 73.72 | 144.91 | | |

**BlueModule: Streptococcus.2**

| OTU_Name | Genus | Abundance% | Prevalence% | -log10(*P*) MM | -log10(*P*) Smo | -log10(*P*) Asth |
|---|---|---|---|---|---|---|
| Streptococcus_4768 | Streptococcus | 11.22 | 100.00 | 61.10 | | |
| Haemophilus_11091 | Haemophilus | 2.75 | 100.00 | 90.16 | | |
| Veillonella_19390 | Veillonella | 1.87 | 100.00 | 48.69 | | |
| Gemella_3258 | Gemella | 1.64 | 100.00 | 97.37 | | |
| Granulicatella_3979 | Granulicatella | 1.40 | 100.00 | 109.38 | | |
| Haemophilus_11056 | Haemophilus | 0.38 | 80.72 | 14.71 | | |
| Pasteurellaceae_11461 | Unknown | 0.28 | 97.35 | 46.26 | | |
| Veillonella_19388 | Veillonella | 0.22 | 99.43 | 48.13 | | |
| Haemophilus_10797 | Haemophilus | 0.22 | 97.54 | 34.24 | | |
| Veillonella_19389 | Veillonella | 0.21 | 99.62 | 57.86 | | |
| Streptococcus_4755 | Streptococcus | 0.03 | 99.62 | 66.34 | | |
| Rothia_13982 | Rothia | 0.20 | 99.05 | 50.22 | | |
| Streptococcus_4687 | Streptococcus | 0.03 | 98.87 | 63.44 | | |
| Lactobacillales_4201 | Unknown | 0.03 | 92.82 | 168.95 | | |
| Streptococcus_7928 | Streptococcus | 0.08 | 97.16 | 156.20 | | |
| Gemella_3389 | Gemella | 0.01 | 87.71 | 150.50 | | |
| Gemella_3253 | Gemella | 0.03 | 94.14 | 149.91 | | |
| Bacillales_3452 | Unknown | 0.02 | 87.15 | 149.80 | | |
| Bacillales_3375 | Unknown | 0.01 | 75.43 | 149.55 | | |
| Granulicatella_3474 | Granulicatella | 0.01 | 74.10 | 142.50 | | |
| Streptococcus_6167 | Streptococcus | 0.02 | 91.68 | 137.98 | | |
| Gemella_3415 | Gemella | 0.02 | 87.71 | 135.65 | | |
| Gemella_3297 | Gemella | 0.01 | 77.13 | 131.99 | | |

**MagentaModule: Streptococcus.1**

| OTU_Name | Genus | Abundance% | Prevalence% | -log10(*P*) MM | -log10(*P*) Smo | -log10(*P*) Asth |
|---|---|---|---|---|---|---|
| Firmicutes_6640 | Unidentified_Firmicutes | 11.79 | 100.00 | 98.27 | | |
| Streptococcus_20297 | Streptococcus | 6.92 | 100.00 | 241.97 | 5.52 | |
| Streptococcus_7798 | Streptococcus | 0.15 | 100.00 | 42.85 | | |
| Streptococcus_20338 | Streptococcus | 0.14 | 99.62 | 140.61 | | |
| Streptococcus_20302 | Streptococcus | 0.09 | 99.81 | 265.10 | 4.47 | |
| Streptococcus_6916 | Streptococcus | 0.05 | 99.24 | 146.74 | | |
| Streptococcus_6815 | Streptococcus | 0.04 | 94.52 | 88.55 | | |
| Streptococcus_5304 | Streptococcus | 0.03 | 99.62 | 148.90 | | |
| Streptococcus_20310 | Streptococcus | 0.02 | 96.98 | 205.36 | | |
| Rothia_13888 | Rothia | 0.02 | 90.93 | 26.77 | | |
| Streptococcus_8021 | Streptococcus | 0.01 | 96.41 | 164.18 | | |
| Streptococcus_20305 | Streptococcus | 0.02 | 95.27 | 152.52 | | |
| Streptococcus_20337 | Streptococcus | 0.01 | 95.09 | 155.30 | | |
| Streptococcus_7171 | Streptococcus | 0.01 | 94.14 | 130.37 | | |

**BrownModule: Fusobacteria**

| OTU_Name | Genus | Abundance% | Prevalence% | -log10(*P*) MM | -log10(*P*) Smo | -log10(*P*) Asth |
|---|---|---|---|---|---|---|
| Fusobacterium_7409 | Fusobacterium | 4.40 | 100.00 | 149.82 | 6.27 | |
| Leptotrichia_8995 | Leptotrichia | 0.74 | 95.84 | 120.71 | 7.16 | |
| Prevotella_621 | Prevotella | 0.33 | 93.57 | 78.30 | | |
| Oribacterium_15965 | Oribacterium | 0.29 | 99.05 | 69.50 | | |
| Stomatobaculum_15638 | Stomatobaculum | 0.19 | 90.93 | 77.39 | | |
| Lachnoanaerobaculum_14972 | Lachnoanaerobaculum | 0.17 | 97.54 | 99.28 | | |
| Peptostreptococcus_20146 | Peptostreptococcus | 0.17 | 91.87 | 95.44 | | |
| Prevotella_2498 | Prevotella | 0.10 | 71.83 | 21.89 | | |
| Tannerella_312 | Tannerella | 0.09 | 81.29 | 74.55 | | |
| Prevotella_1736 | Prevotella | 0.06 | 82.04 | 94.58 | | |
| Catonella_9245 | Catonella | 0.05 | 93.57 | 82.14 | | |

| | | | | | | |
|---|---|---|---|---|---|---|
| Clostridiales_8204 | Unknown | 0.02 | 83.18 | 51.25 | | |
| Capnocytophaga_2417 | Capnocytophaga | 0.06 | 82.61 | 35.61 | | 2.55 |
| Leptotrichia_8999 | Leptotrichia | 0.03 | 64.46 | 115.42 | | |
| Prevotella_660 | Prevotella | 0.01 | 55.58 | 93.98 | | |
| Peptostreptococcus_20150 | Peptostreptococcus | 0.01 | 55.58 | 89.53 | | |
| Peptostreptococcus_20149 | Peptostreptococcus | 0.01 | 55.58 | 84.71 | | |
| Leptotrichia_8997 | Leptotrichia | 0.00 | 39.51 | 83.14 | | |

**GreenModule: Neisseria**

| OTU_Name | Genus | Abundance% | Prevalence% | -log10*(P)* MM | -log10*(P)* Smo | -log10*(P)* Asth |
|---|---|---|---|---|---|---|
| Neisseria_10019 | Neisseria | 3.97 | 99.24 | 243.35 | 15.16 | |
| Porphyromonas_11896 | Porphyromonas | 1.13 | 98.11 | 83.71 | | |
| Neisseria_9962 | Neisseria | 0.36 | 94.90 | 119.40 | | |
| Capnocytophaga_2454 | Capnocytophaga | 0.26 | 91.30 | 43.16 | | 4.49 |
| Capnocytophaga_509 | Capnocytophaga | 0.26 | 93.57 | 62.39 | | |
| Neisseria_9888 | Neisseria | 0.18 | 80.91 | 113.68 | | |
| Capnocytophaga_2384 | Capnocytophaga | 0.15 | 75.43 | 36.66 | | 1.61 |
| Bergeyella_430 | Bergeyella | 0.08 | 92.63 | 83.05 | | |
| Haemophilus_11369 | Haemophilus | 0.05 | 74.10 | 206.44 | | |
| Neisseriaceae_10099 | Unknown | 0.04 | 82.04 | 201.65 | | |
| Neisseria_10209 | Neisseria | 0.03 | 85.44 | 74.46 | | |
| Neisseria_10178 | Neisseria | 0.03 | 79.40 | 219.70 | | |
| Neisseria_10020 | Neisseria | 0.04 | 75.05 | 230.93 | 12.58 | |
| Neisseria_14366 | Neisseria | 0.03 | 71.27 | 209.30 | | |
| Neisseria_10062 | Neisseria | 0.02 | 69.00 | 202.89 | | |
| Neisseria_9813 | Neisseria | 0.01 | 57.47 | 153.62 | | |
| Neisseria_10095 | Neisseria | 0.01 | 56.33 | 153.60 | | |
| Neisseria_9715 | Neisseria | 0.00 | 52.17 | 128.68 | | |

**BlackModule: Prevotella.2**

| OTU_Name | Genus | Abundance% | Prevalence% | -log10*(P)* MM | -log10*(P)* Smo | -log10*(P)* Asth |
|---|---|---|---|---|---|---|
| Parvimonas_8503 | Parvimonas | 0.16 | 96.98 | 78.19 | | |
| Streptococcus_28 | Streptococcus | 0.14 | 98.30 | 83.83 | | |
| Porphyromonas_261 | Porphyromonas | 0.13 | 79.40 | 84.33 | | |
| Prevotella_1791 | Prevotella | 0.06 | 87.33 | 92.42 | | |
| Prevotella_2177 | Prevotella | 0.05 | 86.20 | 79.98 | | |
| Filifactor_8388 | Filifactor | 0.05 | 56.52 | 75.32 | | |
| Prevotella_1014 | Prevotella | 0.04 | 50.09 | 46.45 | | |
| Clostridiales_8242 | Unknown | 0.04 | 74.48 | 110.42 | 3.85 | |
| Fusobacterium_7664 | Fusobacterium | 0.03 | 12.10 | 11.89 | | |
| Veillonellaceae_19943 | Unknown | 0.02 | 65.60 | 95.50 | | |
| Paludibacter_2300 | Paludibacter | 0.01 | 71.27 | 31.48 | | |
| Mycoplasma_12299 | Mycoplasma | 0.01 | 58.60 | 58.47 | | |
| Tannerella_307 | Tannerella | 0.01 | 52.17 | 72.49 | | |
| Synergistaceae_8518 | Unknown | 0.00 | 38.00 | 68.89 | | |

**CyanModule: Veillonella**

| OTU_Name | Genus | Abundance% | Prevalence% | -log10*(P)* MM | -log10*(P)* Smo | -log10*(P)* Asth |
|---|---|---|---|---|---|---|
| Veillonella_18212 | Veillonella | 0.37 | 71.83 | 131.39 | 6.76 | |
| Veillonella_19620 | Veillonella | 0.00 | 34.78 | 97.93 | 2.22 | |
| Firmicutes_18412 | Unknown | 0.00 | 23.06 | 80.79 | | |
| Firmicutes_18411 | Unknown | 0.00 | 22.68 | 78.64 | | |
| Veillonella_18239 | Veillonella | 0.00 | 14.74 | 47.61 | | |
| Firmicutes_18226 | Unknown | 0.00 | 16.45 | 57.42 | | |
| Veillonella_18283 | Veillonella | 0.00 | 12.10 | 40.64 | | |
| Clostridiales_18253 | Unknown | 0.00 | 15.50 | 50.33 | | |
| Atopobium_5807 | Atopobium | 0.00 | 16.07 | 45.32 | | |
| Veillonella_18222 | Veillonella | 0.00 | 12.48 | 41.61 | | |
| Veillonella_19619 | Veillonella | 0.00 | 19.66 | 31.64 | | |

| | | | | |
|---|---|---|---|---|
| Veillonella_19463 | Veillonella | 0.00 | 18.15 | 41.44 |
| Veillonella_19991 | Veillonella | 0.00 | 17.58 | 40.93 |
| Veillonella_18289 | Veillonella | 0.00 | 12.67 | 41.53 |

**PurpleModule: Prevotella.3**

| **OTU_Name** | **Genus** | **Abundance%** | **Prevalence%** | **-log10*(P)* MM** | **-log10*(P)* Smo** | **-log10*(P)* Asth** |
|---|---|---|---|---|---|---|
| Prevotella_2890 | Prevotella | 0.21 | 84.31 | 117.63 | | |
| RF9_12250 | | 0.01 | 18.71 | 22.85 | | |
| Prevotella_1442 | Prevotella | 0.00 | 26.09 | 67.50 | | |
| Prevotella_2912 | Prevotella | 0.00 | 25.52 | 65.56 | | |
| Clostridiales_8287 | Unknown | 0.00 | 31.57 | 33.68 | | |
| Capnocytophaga_499 | Capnocytophaga | 0.00 | 15.88 | 15.09 | | |
| Prevotella_2816 | Prevotella | 0.00 | 17.96 | 44.24 | | |
| Prevotella_2891 | Prevotella | 0.00 | 18.53 | 49.62 | | |
| Prevotella_1431 | Prevotella | 0.00 | 18.90 | 47.71 | | |
| Prevotella_2146 | Prevotella | 0.00 | 16.64 | 45.84 | | |
| Prevotella_773 | Prevotella | 0.00 | 21.93 | 30.55 | | |
| Prevotella_2732 | Prevotella | 0.00 | 16.45 | 41.06 | | |
| Prevotella_2913 | Prevotella | 0.00 | 10.59 | 32.28 | | |

**TanModule: Indeterminate**

| **OTU_Name** | **Genus** | **Abundance%** | **Prevalence%** | **-log10*(P)* MM** | **-log10*(P)* Smo** | **-log10*(P)* Asth** |
|---|---|---|---|---|---|---|
| Candidatedivision_SR1_12158 | | 0.11 | 64.46 | 73.85 | | |
| Prevotella_822 | Prevotella | 0.07 | 71.08 | 123.87 | 3.27 | |
| RF9_12187 | | 0.01 | 34.59 | 35.69 | | |
| Candidatedivision_SR1_12141 | | 0.01 | 24.39 | 41.68 | | |
| Treponema_11646 | Treponema | 0.01 | 33.46 | 46.11 | | |
| Prevotella_1354 | Prevotella | 0.00 | 29.11 | 75.12 | 1.89 | |
| Prevotella_809 | Prevotella | 0.00 | 23.82 | 66.19 | | |
| Treponema_11722 | Treponema | 0.00 | 14.56 | 29.96 | | |
| Prevotella_848 | Prevotella | 0.00 | 16.64 | 48.81 | | |

| | | | | |
|---|---|---|---|---|
| Candidatedivision_SR1_12142 | | 0.00 | 9.83 | 20.74 |
| Prevotella_2653 | Prevotella | 0.00 | 17.58 | 52.94 |
| Prevotella_640 | Prevotella | 0.00 | 15.12 | 43.37 |
| Prevotella_858 | Prevotella | 0.00 | 15.12 | 44.50 |

**SalmonModule: Porphymonas**

| OTU_Name | Genus | Abundance% | Prevalence% | -log10*(P)* MM | -log10*(P)* Smo | -log10*(P)* Asth |
|---|---|---|---|---|---|---|
| Porphyromonas_598 | Porphyromonas | 0.19 | 16.45 | 22.20 | | |
| Prevotella_1923 | Prevotella | 0.00 | 15.31 | 34.53 | 2.52 | |
| Bacteroides_612 | Bacteroides | 0.00 | 0.38 | 1.88 | | |
| Prevotella_992 | Prevotella | 0.00 | 10.78 | 33.09 | | |
| Prevotella_1003 | Prevotella | 0.00 | 0.76 | 2.40 | | |
| Lachnospiraceae_15543 | Unknown | 0.00 | 3.97 | 8.30 | | |
| Porphyromonas_262 | Porphyromonas | 0.00 | 11.34 | 28.07 | | |
| Porphyromonas_588 | Porphyromonas | 0.00 | 1.32 | 4.83 | | |
| Desulfovibrio_12642 | Desulfovibrio | 0.00 | 1.89 | 5.07 | | |
| Akkermansia_12380 | Akkermansia | 0.00 | 0.38 | 1.95 | | |
| Porphyromonas_2279 | Porphyromonas | 0.00 | 7.75 | 16.86 | | |
| Porphyromonas_11948 | Porphyromonas | 0.00 | 7.37 | 16.33 | | |
| Porphyromonas_2284 | Porphyromonas | 0.00 | 6.24 | 18.04 | | |
| Prevotella_1016 | Prevotella | 0.00 | 5.67 | 14.79 | | |
| Prevotella_1008 | Prevotella | 0.00 | 5.48 | 16.29 | | |
| Prevotella_1319 | Prevotella | 0.00 | 5.48 | 15.80 | | |

**PinkModule: Bifidobacteria**

| OTU_Name | Genus | Abundance% | Prevalence% | -log10*(P)* MM | -log10*(P)* Smo | -log10*(P)* Asth |
|---|---|---|---|---|---|---|
| Bifidobacterium_13861 | Bifidobacterium | 0.08 | 47.07 | 42.25 | 14.06 | |
| Streptococcus_4578 | Streptococcus | 0.02 | 60.49 | 36.18 | | |
| Lactobacillus_4348 | Lactobacillus | 0.01 | 11.34 | 24.34 | | |
| Streptococcus_226 | Streptococcus | 0.01 | 16.82 | 14.61 | | |
| Lactobacillus_4496 | Lactobacillus | 0.01 | 17.77 | 29.17 | | |

| | | | | | |
|---|---|---|---|---|---|
| Lactobacillus_4421 | Lactobacillus | 0.01 | 18.90 | 32.18 | |
| Bifidobacteriaceae_13837 | Unknown | 0.01 | 35.92 | 33.08 | |
| Prevotella_1947 | Prevotella | 0.01 | 7.37 | 5.79 | |
| Lactobacillus_4457 | Lactobacillus | 0.00 | 6.24 | 14.67 | |
| Streptococcus_4601 | Streptococcus | 0.00 | 29.11 | 33.12 | |
| Bifidobacteriaceae_13749 | Unknown | 0.00 | 18.90 | 29.72 | |
| Veillonella_20045 | Veillonella | 0.00 | 18.71 | 15.40 | |
| Bifidobacterium_13401 | Bifidobacterium | 0.00 | 16.45 | 37.23 | 10.40 |
| Bifidobacterium_13785 | Bifidobacterium | 0.00 | 14.56 | 31.50 | |

**MidnightBlueModule: Peptococcus**

| OTU_Name | Genus | Abundance% | Prevalence% | -log10*(P)* MM | -log10*(P)* Smo | -log10*(P)* Asth |
|---|---|---|---|---|---|---|
| Peptococcus_9344 | Peptococcus | 0.16 | 88.09 | 202.84 | 4.14 | |
| Peptococcus_9267 | Peptococcus | 0.01 | 47.07 | 133.05 | 4.35 | |
| Firmicutes_16800 | Unknown | 0.00 | 32.51 | 90.04 | | |
| Peptococcus_9274 | Peptococcus | 0.00 | 29.11 | 88.23 | | |
| Peptococcus_9298 | Peptococcus | 0.00 | 27.79 | 81.17 | | |
| Peptococcus_9351 | Peptococcus | 0.00 | 17.01 | 48.58 | | |
| Veillonellaceae_18178 | Unknown | 0.00 | 24.39 | 55.69 | | |
| Peptococcus_9291 | Peptococcus | 0.00 | 18.53 | 45.76 | | |
| Veillonella_18183 | Veillonella | 0.00 | 21.36 | 59.61 | | |
| Peptococcus_9371 | Peptococcus | 0.00 | 17.77 | 45.34 | | |
| Peptococcus_9299 | Peptococcus | 0.00 | 20.60 | 52.84 | | |
| Streptococcus_5498 | Streptococcus | 0.00 | 20.23 | 51.36 | | |
| Veillonella_18513 | Veillonella | 0.00 | 18.71 | 46.91 | | |
| Peptococcus_9337 | Peptococcus | 0.00 | 17.77 | 50.04 | | |

**GreenYellowModule: Contaminants**

| OTU_Name | Genus | Abundance% | Prevalence% | -log10*(P)* MM | -log10*(P)* Smo | -log10*(P)* Asth |
|---|---|---|---|---|---|---|
| Haemophilus_11389 | Haemophilus | 0.01 | 50.47 | 109.82 | 2.75 | |
| Neisseria_9743 | Neisseria | 0.00 | 32.14 | 76.18 | 3.21 | |

| | | | | |
|---|---|---|---|---|
| Herbaspirillum_10515 | Herbaspirillum | 0.00 | 23.06 | 71.10 |
| Herbaspirillum_10738 | Herbaspirillum | 0.00 | 13.99 | 47.11 |
| Haemophilus_11362 | Haemophilus | 0.00 | 14.56 | 29.66 |
| Streptococcus_5394 | Streptococcus | 0.00 | 15.69 | 43.41 |
| Neisseria_9863 | Neisseria | 0.00 | 8.88 | 26.43 |
| Haemophilus_11176 | Haemophilus | 0.00 | 13.42 | 36.60 |
| Pseudomonas_12457 | Pseudomonas | 0.00 | 10.59 | 32.97 |
| Veillonellaceae_16762 | Unknown | 0.00 | 11.53 | 34.35 |
| Actinomyces_13512 | Actinomyces | 0.00 | 15.31 | 19.99 |
| Firmicutes_5407 | Unknown | 0.00 | 11.72 | 36.36 |
| Pseudomonas_12488 | Pseudomonas | 0.00 | 10.78 | 35.41 |

**GreyModule: Unconnected**

| OTU_Name | Genus | Abundance% | Prevalence% |
|---|---|---|---|
| Vibrio_11550 | Vibrio | 0.00 | 2.27 |
| Lactobacillales_3493 | Unknown | 0.00 | 0.76 |
| Corynebacterium_14237 | Corynebacterium | 0.00 | 2.08 |
| Comamonadaceae_10394 | Unknown | 0.00 | 1.89 |
| Escherichia_Shigella_9556 | Escherichia_Shigella | 0.00 | 1.32 |
| Corynebacterium_14199 | Corynebacterium | 0.00 | 2.65 |
| Pseudomonadaceae_12534 | Unknown | 0.00 | 1.89 |
| Burkholderia_10482 | Burkholderia | 0.00 | 1.70 |
| Enterobacter_9544 | Enterobacter | 0.00 | 1.70 |
| vadinBB60_12276 | Unknown | 0.00 | 0.38 |
| Haemophilus_10769 | Haemophilus | 0.00 | 2.08 |
| Vibrio_11520 | Vibrio | 0.00 | 1.89 |
| Pasteurellaceae_11449 | Unknown | 0.00 | 1.51 |

Supplementary Table 5a. Individual OTUs increased in smokers

| OUT_ID | Genus | Fold_change | -log10(*P*) | Abundance | Abundance % | Prevalence % | Change | Increase |
|---|---|---|---|---|---|---|---|---|
| Firmicutes_6640 | Unidentified_Firmicutes | 1.20 | 10.74 | 5146895 | 11.79% | 100.00 | 11846325 | 6,699,430 |
| Veillonella_19412 | Veillonella | 0.70 | 4.60 | 4763249 | 10.91% | 100.00 | 7723612 | 2,960,363 |
| Streptococcus_20297 | Streptococcus | 1.46 | 27.96 | 3019924 | 6.92% | 100.00 | 8285093 | 5,265,169 |
| Rothia_14025 | Rothia | 1.93 | 4.87 | 69239 | 0.16% | 76.75 | 263551 | 194,312 |
| Streptococcus_28 | Streptococcus | 1.19 | 3.71 | 62335 | 0.14% | 98.30 | 142258 | 79,923 |
| Streptococcus_20338 | Streptococcus | 1.13 | 12.02 | 61991 | 0.14% | 99.62 | 135873 | 73,882 |
| Streptococcus_20302 | Streptococcus | 1.40 | 25.63 | 39744 | 0.09% | 99.81 | 104758 | 65,014 |
| Bifidobacterium_13861 | Bifidobacterium | 3.17 | 5.14 | 36752 | 0.08% | 47.07 | 330810 | 294,058 |
| Prevotella_1791 | Prevotella | 1.35 | 3.72 | 25413 | 0.06% | 87.33 | 65005 | 39,592 |
| Streptococcus_6916 | Streptococcus | 1.05 | 6.00 | 22255 | 0.05% | 99.24 | 46124 | 23,869 |
| Streptococcus_6617 | Streptococcus | 0.69 | 4.48 | 14951 | 0.03% | 97.73 | 24172 | 9,221 |
| Streptococcus_5304 | Streptococcus | 1.14 | 14.57 | 14557 | 0.03% | 99.62 | 32024 | 17,467 |
| Actinomyces_13500 | Actinomyces | 0.92 | 4.63 | 13457 | 0.03% | 96.41 | 25494 | 12,037 |
| Streptococcus_4578 | Streptococcus | 2.66 | 7.38 | 10289 | 0.02% | 60.49 | 64849 | 54,560 |
| Bifidobacteriaceae_13811 | Unknown | 2.93 | 5.30 | 10032 | 0.02% | 42.53 | 76595 | 66,563 |
| Streptococcus_20310 | Streptococcus | 1.16 | 12.62 | 9713 | 0.02% | 96.98 | 21691 | 11,978 |
| Streptococcus_4835 | Streptococcus | 0.65 | 3.74 | 9083 | 0.02% | 95.46 | 14228 | 5,145 |
| Rothia_13888 | Rothia | 1.98 | 14.51 | 8711 | 0.02% | 90.93 | 34480 | 25,769 |
| Streptococcus_20305 | Streptococcus | 1.26 | 12.49 | 6569 | 0.02% | 95.27 | 15743 | 9,174 |
| Leptotrichia_8836 | Leptotrichia | 1.26 | 3.41 | 6223 | 0.01% | 75.61 | 14873 | 8,650 |
| Lactobacillus_4348 | Lactobacillus | 4.46 | 3.75 | 5290 | 0.01% | 11.34 | 116661 | 111,371 |
| Streptococcus_8021 | Streptococcus | 1.26 | 13.46 | 5262 | 0.01% | 96.41 | 12611 | 7,349 |
| Streptococcus_20337 | Streptococcus | 1.57 | 20.46 | 4872 | 0.01% | 95.09 | 14452 | 9,580 |
| Streptococcus_7171 | Streptococcus | 1.43 | 17.05 | 4860 | 0.01% | 94.14 | 13098 | 8,238 |
| Streptococcus_6449 | Streptococcus | 0.82 | 6.69 | 4521 | 0.01% | 91.68 | 7977 | 3,456 |
| Streptococcus_226 | Streptococcus | 4.05 | 3.67 | 4212 | 0.01% | 16.82 | 69867 | 65,655 |
| Lactobacillus_4496 | Lactobacillus | 4.39 | 6.74 | 3357 | 0.01% | 17.77 | 70554 | 67,197 |
| Streptococcus_7797 | Streptococcus | 1.08 | 7.13 | 3041 | 0.01% | 88.09 | 6428 | 3,387 |

| | | | | | | | | |
|---|---|---|---|---|---|---|---|---|
| Bifidobacteriaceae_13837 | Unknown | 2.74 | 4.90 | 2741 | 0.01% | 35.92 | 18369 | 15,628 |
| Bifidobacteriaceae_13747 | Unknown | 3.44 | 12.31 | 2256 | 0.01% | 42.34 | 24439 | 22,183 |
| Streptococcus_5317 | Streptococcus | 1.09 | 6.44 | 1767 | 0.00% | 80.72 | 3764 | 1,997 |
| Cryptobacterium_12323 | Cryptobacterium | 3.02 | 4.14 | 1716 | 0.00% | 23.44 | 13952 | 12,236 |
| Streptococcus_4601 | Streptococcus | 2.96 | 5.92 | 1455 | 0.00% | 29.11 | 11327 | 9,872 |
| Bifidobacteriaceae_13749 | Unknown | 2.73 | 3.21 | 1429 | 0.00% | 18.90 | 9474 | 8,045 |
| Streptococcus_7277 | Streptococcus | 0.89 | 4.48 | 1370 | 0.00% | 71.64 | 2542 | 1,172 |
| Streptococcus_20306 | Streptococcus | 1.40 | 6.20 | 1355 | 0.00% | 70.70 | 3575 | 2,220 |
| Streptococcus_20342 | Streptococcus | 1.44 | 10.30 | 1300 | 0.00% | 73.16 | 3539 | 2,239 |
| Streptococcus_7114 | Streptococcus | 1.27 | 8.89 | 1289 | 0.00% | 67.49 | 3103 | 1,814 |
| Streptococcus_20298 | Streptococcus | 1.12 | 8.56 | 1168 | 0.00% | 72.02 | 2546 | 1,378 |
| Streptococcus_7048 | Streptococcus | 1.65 | 15.23 | 1104 | 0.00% | 71.64 | 3459 | 2,355 |
| Streptococcus_20289 | Streptococcus | 1.42 | 8.54 | 971 | 0.00% | 68.05 | 2603 | 1,632 |
| Streptococcus_20326 | Streptococcus | 1.44 | 10.46 | 946 | 0.00% | 65.78 | 2571 | 1,625 |
| Streptococcus_7148 | Streptococcus | 1.08 | 5.62 | 852 | 0.00% | 65.41 | 1807 | 955 |
| Streptococcus_20335 | Streptococcus | 1.03 | 3.96 | 850 | 0.00% | 60.68 | 1738 | 888 |
| Dialister_19867 | Dialister | 3.88 | 3.03 | 816 | 0.00% | 10.21 | 12009 | 11,193 |
| Streptococcus_7230 | Streptococcus | 1.13 | 4.85 | 746 | 0.00% | 59.17 | 1634 | 888 |
| Streptococcus_7017 | Streptococcus | 1.05 | 4.60 | 746 | 0.00% | 62.95 | 1544 | 798 |
| Streptococcus_7180 | Streptococcus | 1.04 | 4.83 | 743 | 0.00% | 62.00 | 1530 | 787 |
| Streptococcus_7143 | Streptococcus | 1.42 | 9.75 | 663 | 0.00% | 58.22 | 1777 | 1,114 |
| Streptococcus_8017 | Streptococcus | 1.11 | 5.12 | 598 | 0.00% | 53.69 | 1290 | 692 |
| Streptococcus_7139 | Streptococcus | 1.18 | 6.20 | 575 | 0.00% | 55.58 | 1300 | 725 |
| Bifidobacterium_13401 | Bifidobacterium | 3.13 | 4.05 | 528 | 0.00% | 16.45 | 4627 | 4,099 |
| Streptococcus_7865 | Streptococcus | 2.72 | 6.69 | 517 | 0.00% | 25.71 | 3405 | 2,888 |
| Bifidobacterium_13785 | Bifidobacterium | 3.13 | 3.61 | 514 | 0.00% | 14.56 | 4512 | 3,998 |
| Streptococcus_5678 | Streptococcus | 1.11 | 3.35 | 498 | 0.00% | 44.42 | 1072 | 574 |
| Streptococcus_8030 | Streptococcus | 1.14 | 4.45 | 485 | 0.00% | 47.45 | 1070 | 585 |
| Streptococcus_7159 | Streptococcus | 1.19 | 4.48 | 455 | 0.00% | 44.80 | 1038 | 583 |
| Streptococcus_8008 | Streptococcus | 0.97 | 3.64 | 454 | 0.00% | 46.69 | 892 | 438 |
| Rothia_13334 | Rothia | 1.65 | 3.01 | 449 | 0.00% | 27.79 | 1411 | 962 |

| | | | | | | | | |
|---|---|---|---|---|---|---|---|---|
| Streptococcus_4641 | Streptococcus | 1.05 | 3.48 | 440 | 0.00% | 47.45 | 910 | 470 |
| Howardella_9190 | Howardella | 3.15 | 5.98 | 413 | 0.00% | 18.53 | 3654 | 3,241 |
| Streptococcus_7225 | Streptococcus | 1.33 | 4.45 | 372 | 0.00% | 41.02 | 936 | 564 |
| Streptococcus_7813 | Streptococcus | 1.15 | 3.56 | 296 | 0.00% | 35.16 | 655 | 359 |
| Streptococcus_7760 | Streptococcus | 1.46 | 6.11 | 290 | 0.00% | 36.67 | 797 | 507 |
| Actinomyces_13458 | Actinomyces | 1.59 | 3.36 | 275 | 0.00% | 26.47 | 826 | 551 |

#### Supplementary Table 5b. OTUs decreased in smokers

| OUT_ID | Genus | Fold_change | -log10(*P*) | Abundance | Abundance % | Prevalence % | Change | Increase |
|---|---|---|---|---|---|---|---|---|
| Fusobacterium_7409 | Fusobacterium | -1.30 | 7.01 | 1921431 | 3.97% | 100.00 | 781164 | -1,140,267 |
| Neisseria_10019 | Neisseria | -3.32 | 23.88 | 1735113 | 2.75% | 99.24 | 173906 | -1,561,207 |
| Haemophilus_11091 | Haemophilus | -1.58 | 10.67 | 1199062 | 1.87% | 100.00 | 401269 | -797,793 |
| Veillonella_19390 | Veillonella | -0.88 | 5.09 | 817686 | 1.64% | 100.00 | 444007 | -373,679 |
| Gemella_3258 | Gemella | -1.08 | 5.78 | 714904 | 1.13% | 100.00 | 337850 | -377,054 |
| Porphyromonas_11896 | Porphyromonas | -3.04 | 18.95 | 491972 | 0.79% | 98.11 | 59970 | -432,002 |
| Prevotella_1969 | Prevotella | -1.13 | 3.67 | 346641 | 0.74% | 98.11 | 158588 | -188,053 |
| Leptotrichia_8995 | Leptotrichia | -2.31 | 10.74 | 324978 | 0.49% | 95.84 | 65668 | -259,310 |
| Leptotrichia_8876 | Leptotrichia | -2.50 | 4.75 | 215430 | 0.36% | 68.43 | 38202 | -177,228 |
| Neisseria_9962 | Neisseria | -2.91 | 23.19 | 158791 | 0.33% | 94.90 | 21191 | -137,600 |
| Prevotella_621 | Prevotella | -2.88 | 17.71 | 144656 | 0.29% | 93.57 | 19602 | -125,054 |
| Oribacterium_15965 | Oribacterium | -1.10 | 6.02 | 124840 | 0.28% | 99.05 | 58441 | -66,399 |
| Pasteurellaceae_11461 | Unknown | -2.14 | 10.61 | 123462 | 0.26% | 97.35 | 28037 | -95,425 |
| Capnocytophaga_2454 | Capnocytophaga | -3.24 | 15.96 | 113957 | 0.26% | 91.30 | 12083 | -101,874 |
| Capnocytophaga_509 | Capnocytophaga | -3.71 | 22.58 | 113027 | 0.22% | 93.57 | 8622 | -104,405 |
| Veillonella_19388 | Veillonella | -0.89 | 3.57 | 97522 | 0.22% | 99.43 | 52686 | -44,836 |
| Haemophilus_10797 | Haemophilus | -1.37 | 4.89 | 96360 | 0.19% | 97.54 | 37377 | -58,983 |
| Stomatobaculum_15638 | Stomatobaculum | -1.94 | 9.03 | 81904 | 0.18% | 90.93 | 21419 | -60,485 |
| Neisseria_9888 | Neisseria | -3.68 | 18.95 | 76462 | 0.17% | 80.91 | 5961 | -70,501 |
| Lachnoanaerobaculum_14972 | Lachnoanaerobaculum | -1.08 | 4.37 | 75543 | 0.17% | 97.54 | 35757 | -39,786 |

| | | | | | | | | |
|---|---|---|---|---|---|---|---|---|
| Peptostreptococcus_20146 | Peptostreptococcus | -1.83 | 8.30 | 74697 | 0.16% | 91.87 | 20946 | -53,751 |
| Peptococcus_9344 | Peptococcus | -1.93 | 6.38 | 70217 | 0.15% | 88.09 | 18468 | -51,749 |
| Capnocytophaga_2384 | Capnocytophaga | -3.82 | 15.74 | 64767 | 0.11% | 75.43 | 4589 | -60,178 |
| Candidate_division_SR1_12158 | Unknown | -3.67 | 10.38 | 49511 | 0.09% | 64.46 | 3902 | -45,609 |
| Tannerella_312 | Tannerella | -3.01 | 11.94 | 38759 | 0.08% | 81.29 | 4805 | -33,954 |
| Haemophilus_10843 | Haemophilus | -1.48 | 7.66 | 35829 | 0.08% | 95.84 | 12819 | -23,010 |
| Prevotella_2105 | Prevotella | -3.39 | 4.92 | 33863 | 0.08% | 40.45 | 3229 | -30,634 |
| Bergeyella_430 | Bergeyella | -2.04 | 10.83 | 32875 | 0.07% | 92.63 | 7998 | -24,877 |
| Campylobacter_8553 | Campylobacter | -0.94 | 4.51 | 29816 | 0.07% | 97.73 | 15555 | -14,261 |
| Prevotella_822 | Prevotella | -4.06 | 16.99 | 29280 | 0.06% | 71.08 | 1761 | -27,519 |
| Prevotella_1736 | Prevotella | -1.55 | 4.73 | 28316 | 0.05% | 82.04 | 9652 | -18,664 |
| Haemophilus_11369 | Haemophilus | -3.29 | 15.67 | 21647 | 0.05% | 74.10 | 2217 | -19,430 |
| Catonella_9245 | Catonella | -1.16 | 5.58 | 21066 | 0.04% | 93.57 | 9451 | -11,615 |
| Neisseriaceae_10099 | Unknown | -2.05 | 7.98 | 19372 | 0.04% | 82.04 | 4680 | -14,692 |
| Neisseria_10020 | Neisseria | -3.24 | 17.19 | 16590 | 0.03% | 75.05 | 1757 | -14,833 |
| Neisseria_10178 | Neisseria | -3.24 | 18.27 | 15187 | 0.03% | 79.40 | 1609 | -13,578 |
| Neisseria_14366 | Neisseria | -3.54 | 18.01 | 14399 | 0.03% | 71.27 | 1237 | -13,162 |
| Neisseria_10209 | Neisseria | -2.87 | 19.57 | 13604 | 0.03% | 85.44 | 1865 | -11,739 |
| Johnsonella_16130 | Johnsonella | -2.64 | 6.77 | 12779 | 0.03% | 59.17 | 2055 | -10,724 |
| Leptotrichia_8999 | Leptotrichia | -2.33 | 6.65 | 12420 | 0.02% | 64.46 | 2472 | -9,948 |
| Clostridiales_8204 | Unknown | -1.94 | 11.28 | 9268 | 0.02% | 83.18 | 2408 | -6,860 |
| Bacillales_3452 | Unknown | -0.85 | 3.54 | 8582 | 0.02% | 87.15 | 4758 | -3,824 |
| Leptotrichia_9109 | Leptotrichia | -4.06 | 4.48 | 7973 | 0.02% | 22.68 | 479 | -7,494 |
| Neisseria_10062 | Neisseria | -3.21 | 15.65 | 6895 | 0.01% | 69.00 | 747 | -6,148 |
| RF9_12187 | Unknown | -3.76 | 6.20 | 6506 | 0.01% | 34.59 | 479 | -6,027 |
| Neisseriaceae_9657 | Unknown | -1.70 | 4.89 | 5010 | 0.01% | 73.16 | 1538 | -3,472 |
| Leptotrichia_9129 | Leptotrichia | -1.56 | 3.72 | 4896 | 0.01% | 62.00 | 1660 | -3,236 |
| Haemophilus_10776 | Haemophilus | -1.73 | 5.74 | 4866 | 0.01% | 72.02 | 1466 | -3,400 |
| Aerococcaceae_4310 | Unknown | -1.94 | 4.48 | 4763 | 0.01% | 58.98 | 1241 | -3,522 |
| Clostridiales_16964 | Unknown | -0.78 | 3.09 | 4448 | 0.01% | 82.99 | 2588 | -1,860 |
| Neisseria_10216 | Neisseria | -2.44 | 5.33 | 4315 | 0.01% | 49.72 | 794 | -3,521 |

| | | | | | | | | |
|---|---|---|---|---|---|---|---|---|
| Prevotella_660 | Prevotella | -2.18 | 6.04 | 4122 | 0.01% | 55.58 | 912 | -3,210 |
| Peptostreptococcus_20149 | Peptostreptococcus | -1.56 | 3.30 | 3580 | 0.01% | 55.58 | 1212 | -2,368 |
| Neisseria_9645 | Neisseria | -3.31 | 11.27 | 3472 | 0.01% | 54.82 | 350 | -3,122 |
| Alysiella_9616 | Alysiella | -2.97 | 6.14 | 3446 | 0.01% | 40.26 | 439 | -3,007 |
| Prevotella_1378 | Prevotella | -2.49 | 7.20 | 3338 | 0.01% | 52.93 | 594 | -2,744 |
| Clostridiales_16966 | Unknown | -2.03 | 9.31 | 3139 | 0.01% | 70.32 | 768 | -2,371 |
| Peptostreptococcus_20150 | Peptostreptococcus | -2.45 | 7.85 | 2935 | 0.01% | 55.58 | 539 | -2,396 |
| Lautropia_10617 | Lautropia | -2.09 | 6.94 | 2886 | 0.01% | 64.84 | 677 | -2,209 |
| Neisseria_9813 | Neisseria | -3.03 | 11.88 | 2687 | 0.01% | 57.47 | 329 | -2,358 |
| Firmicutes_18478 | Unknown | -0.98 | 3.39 | 2628 | 0.01% | 75.80 | 1333 | -1,295 |
| Neisseria_10095 | Neisseria | -2.80 | 9.95 | 2618 | 0.01% | 56.33 | 376 | -2,242 |
| Peptococcus_9267 | Peptococcus | -1.98 | 4.24 | 2396 | 0.01% | 47.07 | 606 | -1,790 |
| Neisseria_9959 | Neisseria | -2.21 | 5.93 | 2247 | 0.01% | 50.47 | 486 | -1,761 |
| Neisseria_10028 | Neisseria | -2.60 | 7.84 | 2246 | 0.00% | 51.80 | 370 | -1,876 |
| Neisseria_9715 | Neisseria | -2.24 | 6.32 | 2063 | 0.00% | 52.17 | 435 | -1,628 |
| Stomatobaculum_15684 | Stomatobaculum | -2.12 | 6.01 | 2004 | 0.00% | 50.66 | 461 | -1,543 |
| Cardiobacterium_9389 | Cardiobacterium | -1.73 | 6.15 | 1929 | 0.00% | 66.35 | 583 | -1,346 |
| Neisseria_9958 | Neisseria | -2.64 | 8.43 | 1867 | 0.00% | 52.74 | 299 | -1,568 |
| Haemophilus_11492 | Haemophilus | -2.97 | 4.37 | 1830 | 0.00% | 27.41 | 234 | -1,596 |
| Leptotrichia_9102 | Leptotrichia | -2.49 | 3.06 | 1822 | 0.00% | 27.03 | 324 | -1,498 |
| Haemophilus_11011 | Haemophilus | -1.84 | 7.09 | 1804 | 0.00% | 64.46 | 504 | -1,300 |
| Porphyromonas_11969 | Porphyromonas | -2.36 | 5.72 | 1749 | 0.00% | 44.61 | 340 | -1,409 |
| Haemophilus_11405 | Haemophilus | -1.53 | 5.18 | 1729 | 0.00% | 62.76 | 600 | -1,129 |
| Neisseria_10277 | Neisseria | -1.78 | 3.97 | 1712 | 0.00% | 51.04 | 499 | -1,213 |
| Leptotrichia_8997 | Leptotrichia | -2.40 | 5.09 | 1709 | 0.00% | 39.51 | 325 | -1,384 |
| Streptococcus_5402 | Streptococcus | -2.45 | 7.41 | 1577 | 0.00% | 49.15 | 289 | -1,288 |
| Haemophilus_11086 | Haemophilus | -2.23 | 6.04 | 1533 | 0.00% | 47.45 | 326 | -1,207 |
| Haemophilus_11238 | Haemophilus | -1.89 | 3.23 | 1459 | 0.00% | 39.51 | 393 | -1,066 |
| Leptotrichia_9067 | Leptotrichia | -2.24 | 3.72 | 1442 | 0.00% | 34.22 | 306 | -1,136 |
| Gemella_3265 | Gemella | -1.21 | 3.85 | 1355 | 0.00% | 66.92 | 584 | -771 |
| Streptococcus_5404 | Streptococcus | -2.51 | 9.64 | 1332 | 0.00% | 52.93 | 234 | -1,098 |

| | | | | | | | | |
|---|---|---|---|---|---|---|---|---|
| Neisseria_10149 | Neisseria | -2.64 | 7.01 | 1328 | 0.00% | 44.99 | 214 | -1,114 |
| Porphyromonas_11873 | Porphyromonas | -3.01 | 4.78 | 1214 | 0.00% | 27.03 | 151 | -1,063 |
| Haemophilus_11274 | Haemophilus | -1.30 | 3.91 | 1201 | 0.00% | 57.47 | 487 | -714 |
| Prevotella_1348 | Prevotella | -2.40 | 4.54 | 1162 | 0.00% | 35.54 | 221 | -941 |
| Leptotrichia_9006 | Leptotrichia | -2.18 | 4.05 | 1152 | 0.00% | 35.54 | 254 | -898 |
| Neisseria_9833 | Neisseria | -2.79 | 6.77 | 967 | 0.00% | 38.37 | 140 | -827 |
| Blautia_5645 | Blautia | -1.72 | 3.00 | 963 | 0.00% | 39.13 | 293 | -670 |
| Haemophilus_11385 | Haemophilus | -2.04 | 4.83 | 943 | 0.00% | 40.45 | 230 | -713 |
| Prevotella_641 | Prevotella | -2.68 | 3.41 | 923 | 0.00% | 24.20 | 144 | -779 |
| Neisseria_9708 | Neisseria | -2.42 | 4.49 | 900 | 0.00% | 32.14 | 168 | -732 |
| Pasteurellaceae_10837 | Unknown | -1.99 | 4.14 | 899 | 0.00% | 38.19 | 226 | -673 |
| Haemophilus_11104 | Haemophilus | -1.52 | 3.97 | 843 | 0.00% | 46.88 | 294 | -549 |
| Neisseria_9867 | Neisseria | -2.13 | 4.87 | 757 | 0.00% | 39.70 | 173 | -584 |
| Prevotella_2652 | Prevotella | -2.47 | 4.46 | 755 | 0.00% | 29.87 | 136 | -619 |
| Oribacterium_15967 | Oribacterium | -1.42 | 3.61 | 751 | 0.00% | 49.53 | 281 | -470 |
| Prevotella_657 | Prevotella | -2.24 | 3.75 | 745 | 0.00% | 30.43 | 158 | -587 |
| Porphyromonas_11939 | Porphyromonas | -2.35 | 4.17 | 731 | 0.00% | 29.87 | 143 | -588 |
| Neisseria_10024 | Neisseria | -2.16 | 5.11 | 709 | 0.00% | 38.37 | 159 | -550 |
| Neisseria_10221 | Neisseria | -1.78 | 3.46 | 586 | 0.00% | 36.67 | 170 | -416 |
| Streptococcus_6135 | Streptococcus | -1.47 | 3.23 | 447 | 0.00% | 36.48 | 161 | -286 |

Supplementary Table 6a. OTUs increased in asthmatics

| OTU_ID | Genus | Phylum | Fold_change | -log10(*P*) | Abundance | Abundance % | Prevalence % | Change | Increase |
|---|---|---|---|---|---|---|---|---|---|
| Neisseria_10019 | Neisseria | Proteobacteria | 0.96 | 1.45 | 1,371,169 | 4.74% | 99.44 | 2672809 | 1,301,640 |
| Rothia_13982 | Rothia | Actinobacteria | 0.78 | 1.46 | 65582 | 0.23% | 99.15 | 112474 | 46,892 |

Supplementary Table 6b. OTUs decreased in asthmatics

| OUT_ID | Genus | Fold_change | -log10(*P*) | Abundance | Abundance % | Prevalence % | Change | Increase |
|---|---|---|---|---|---|---|---|---|
| Actinomyces_13710 | Actinomyces | -0.63 | 2.01 | 1241550 | 4.29% | 100.00 | 805016 | -436,534 |
| Selenomonas_17559 | Selenomonas | -0.84 | 1.64 | 429036 | 1.48% | 100.00 | 239398 | -189,638 |
| Leptotrichia_8776 | Leptotrichia | -1.05 | 1.83 | 581845 | 2.01% | 99.72 | 281154 | -300,691 |
| Megasphaera_16215 | Megasphaera | -0.76 | 1.61 | 179586 | 0.62% | 99.72 | 105754 | -73,832 |
| Selenomonas_17440 | Selenomonas | -1.76 | 7.16 | 47875 | 0.17% | 99.44 | 14108 | -33,767 |
| Oribacterium_15113 | Oribacterium | -0.69 | 1.36 | 26997 | 0.09% | 97.18 | 16739 | -10,258 |
| Actinomyces_13062 | Actinomyces | -1.29 | 3.44 | 93637 | 0.32% | 95.77 | 38236 | -55,401 |
| Capnocytophaga_2454 | Capnocytophaga | -2.06 | 4.49 | 94930 | 0.33% | 93.24 | 22839 | -72,091 |
| Prevotella_879 | Prevotella | -0.99 | 1.45 | 56933 | 0.20% | 92.96 | 28662 | -28,271 |
| Streptococcus_4754 | Streptococcus | -0.55 | 1.45 | 6265 | 0.02% | 92.68 | 4275 | -1,990 |
| Selenomonas_17724 | Selenomonas | -1.28 | 2.59 | 24494 | 0.08% | 91.83 | 10101 | -14,393 |
| Candidate_division_TM7_12053 | Unknown | -0.93 | 1.48 | 39771 | 0.14% | 91.27 | 20826 | -18,945 |
| Streptococcus_6284 | Streptococcus | -0.74 | 1.83 | 6738 | 0.02% | 90.99 | 4041 | -2,697 |
| Prevotella_2754 | Prevotella | -1.61 | 2.32 | 113240 | 0.39% | 89.30 | 37057 | -76,183 |
| Actinomyces_13534 | Actinomyces | -0.64 | 1.52 | 6955 | 0.02% | 89.01 | 4456 | -2,499 |
| Ruminococcaceae_11808 | Unknown | -1.18 | 2.14 | 24385 | 0.08% | 88.73 | 10771 | -13,614 |
| Prevotella_2177 | Prevotella | -1.07 | 1.48 | 15489 | 0.05% | 85.07 | 7372 | -8,117 |
| Prevotella_2890 | Prevotella | -1.43 | 1.69 | 64968 | 0.22% | 83.38 | 24040 | -40,928 |
| Capnocytophaga_2417 | Capnocytophaga | -1.24 | 2.55 | 22882 | 0.08% | 82.82 | 9662 | -13,220 |
| Tannerella_312 | Tannerella | -1.72 | 2.69 | 29107 | 0.10% | 82.54 | 8860 | -20,247 |
| Actinomyces_13518 | Actinomyces | -0.67 | 1.34 | 2791 | 0.01% | 80.00 | 1758 | -1,033 |
| Ruminococcaceae_11741 | Unknown | -1.16 | 1.40 | 19176 | 0.07% | 78.87 | 8561 | -10,615 |
| Dialister_19884 | Dialister | -1.12 | 1.49 | 6714 | 0.02% | 78.59 | 3096 | -3,618 |
| Capnocytophaga_2384 | Capnocytophaga | -1.39 | 1.61 | 49387 | 0.17% | 78.31 | 18897 | -30,490 |
| Porphyromonas_261 | Porphyromonas | -1.90 | 3.05 | 36872 | 0.13% | 77.18 | 9911 | -26,961 |
| Haemophilus_10776 | Haemophilus | -0.95 | 1.43 | 3449 | 0.01% | 76.06 | 1790 | -1,659 |
| Streptococcus_5343 | Streptococcus | -0.77 | 1.30 | 2154 | 0.01% | 76.06 | 1263 | -891 |
| Butyrivibrio_14576 | Butyrivibrio | -1.36 | 2.02 | 7617 | 0.03% | 73.80 | 2974 | -4,643 |

| | | | | | | | | |
|---|---|---|---|---|---|---|---|---|
| Veillonellaceae_17891 | Unknown | -2.75 | 4.77 | 50349 | 0.17% | 73.52 | 7484 | -42,865 |
| Streptococcus_5636 | Streptococcus | -0.76 | 1.34 | 1494 | 0.01% | 73.52 | 880 | -614 |
| Prevotella_822 | Prevotella | -1.62 | 2.01 | 21047 | 0.07% | 73.24 | 6825 | -14,222 |
| Actinomyces_13193 | Actinomyces | -0.94 | 2.09 | 1226 | 0.00% | 72.39 | 641 | -585 |
| Paludibacter_2300 | Paludibacter | -1.93 | 3.92 | 3191 | 0.01% | 71.55 | 840 | -2,351 |
| Prevotella_2498 | Prevotella | -2.73 | 4.77 | 34158 | 0.12% | 71.27 | 5146 | -29,012 |
| Clostridiales_8242 | Unknown | -1.56 | 2.14 | 9059 | 0.03% | 69.58 | 3080 | -5,979 |
| Veillonellaceae_16318 | Unknown | -0.85 | 1.40 | 1719 | 0.01% | 69.58 | 952 | -767 |
| Veillonella_18212 | Veillonella | -2.12 | 2.21 | 98326 | 0.34% | 69.01 | 22599 | -75,727 |
| Actinomyces_13323 | Actinomyces | -1.51 | 2.96 | 2974 | 0.01% | 68.73 | 1043 | -1,931 |
| Selenomonas_17617 | Selenomonas | -0.92 | 1.33 | 2328 | 0.01% | 68.73 | 1231 | -1,097 |
| Actinomyces_13298 | Actinomyces | -1.55 | 3.13 | 2781 | 0.01% | 67.04 | 953 | -1,828 |
| Candidate_division_SR1_12158 | Unknown | -2.30 | 2.96 | 39969 | 0.14% | 66.76 | 8135 | -31,834 |
| Leptotrichia_8999 | Leptotrichia | -1.17 | 1.30 | 9063 | 0.03% | 66.20 | 4030 | -5,033 |
| Atopobium_5909 | Atopobium | -1.00 | 1.75 | 1678 | 0.01% | 65.92 | 837 | -841 |
| Veillonella_18675 | Veillonella | -1.21 | 2.21 | 1334 | 0.00% | 65.07 | 578 | -756 |
| Actinomyces_13490 | Actinomyces | -1.05 | 1.67 | 1462 | 0.01% | 64.79 | 705 | -757 |
| Actinomyces_13706 | Actinomyces | -0.99 | 1.51 | 1571 | 0.01% | 64.51 | 789 | -782 |
| Veillonellaceae_19943 | Unknown | -1.27 | 1.40 | 5765 | 0.02% | 63.38 | 2385 | -3,380 |
| Leptotrichia_8642 | Leptotrichia | -2.29 | 3.13 | 8815 | 0.03% | 56.06 | 1798 | -7,017 |
| Mycoplasma_12299 | Mycoplasma | -1.94 | 2.41 | 3621 | 0.01% | 54.93 | 945 | -2,676 |
| Candidate_division_TM7_12081 | Unknown | -1.41 | 1.72 | 2061 | 0.01% | 54.93 | 777 | -1,284 |
| Actinomyces_13523 | Actinomyces | -0.89 | 1.45 | 569 | 0.00% | 53.52 | 308 | -261 |
| Veillonella_16337 | Veillonella | -1.50 | 1.83 | 2082 | 0.01% | 52.96 | 736 | -1,346 |
| Prevotella_1045 | Prevotella | -1.44 | 2.13 | 907 | 0.00% | 52.96 | 335 | -572 |
| Actinomyces_13647 | Actinomyces | -1.47 | 2.29 | 1102 | 0.00% | 52.68 | 397 | -705 |
| Filifactor_8388 | Filifactor | -2.63 | 3.27 | 10142 | 0.04% | 51.83 | 1644 | -8,498 |
| Atopobium_5889 | Atopobium | -1.39 | 1.50 | 1759 | 0.01% | 51.55 | 669 | -1,090 |
| Tannerella_307 | Tannerella | -1.67 | 1.94 | 3719 | 0.01% | 49.01 | 1172 | -2,547 |
| Actinomyces_13517 | Actinomyces | -1.03 | 1.48 | 485 | 0.00% | 48.17 | 237 | -248 |
| Prevotella_1014 | Prevotella | -1.83 | 1.72 | 7345 | 0.03% | 47.32 | 2065 | -5,280 |

| | | | | | | | | |
|---|---|---|---|---|---|---|---|---|
| Veillonella_16930 | Veillonella | -1.28 | 1.45 | 798 | 0.00% | 47.32 | 329 | -469 |
| Veillonella_18231 | Veillonella | -1.19 | 1.46 | 619 | 0.00% | 47.04 | 271 | -348 |
| Selenomonas_17387 | Selenomonas | -1.70 | 2.55 | 839 | 0.00% | 46.20 | 258 | -581 |
| Treponema_11662 | Treponema | -2.08 | 2.36 | 2280 | 0.01% | 45.63 | 540 | -1,740 |
| Ruminococcaceae_11796 | Unknown | -1.23 | 1.34 | 803 | 0.00% | 45.63 | 343 | -460 |
| Prevotella_2030 | Prevotella | -1.48 | 1.45 | 1189 | 0.00% | 43.66 | 427 | -762 |
| Selenomonas_17525 | Selenomonas | -1.35 | 1.45 | 684 | 0.00% | 43.10 | 269 | -415 |
| WCHB1_69_2474 | Unknown | -1.67 | 1.48 | 4298 | 0.01% | 42.82 | 1349 | -2,949 |
| Prevotella_2105 | Prevotella | -2.43 | 1.81 | 26280 | 0.09% | 42.54 | 4866 | -21,414 |
| Prevotella_1769 | Prevotella | -1.74 | 2.13 | 872 | 0.00% | 41.69 | 260 | -612 |
| Veillonella_19887 | Veillonella | -1.54 | 1.45 | 847 | 0.00% | 39.15 | 291 | -556 |
| Bifidobacteriaceae_13811 | Unknown | -1.84 | 1.40 | 2836 | 0.01% | 38.87 | 792 | -2,044 |
| Bifidobacteriaceae_13747 | Unknown | -1.50 | 1.45 | 653 | 0.00% | 38.31 | 232 | -421 |
| Prevotella_2133 | Prevotella | -2.12 | 1.50 | 7866 | 0.03% | 35.21 | 1806 | -6,060 |
| Lachnospiraceae_15303 | Unknown | -1.98 | 1.64 | 1153 | 0.00% | 34.37 | 292 | -861 |
| Veillonella_19620 | Veillonella | -2.72 | 2.96 | 1288 | 0.00% | 33.24 | 196 | -1,092 |
| Selenomonas_17406 | Selenomonas | -2.10 | 2.86 | 386 | 0.00% | 32.39 | 90 | -296 |
| Bergeyella_444 | Bergeyella | -3.10 | 2.96 | 1738 | 0.01% | 29.01 | 203 | -1,535 |
| Butyrivibrio_14621 | Butyrivibrio | -1.83 | 1.42 | 564 | 0.00% | 28.73 | 159 | -405 |
| Prevotella_2584 | Prevotella | -3.56 | 3.44 | 1680 | 0.01% | 28.17 | 143 | -1,537 |
| Prevotella_2151 | Prevotella | -2.06 | 1.45 | 833 | 0.00% | 26.48 | 200 | -633 |
| Treponema_11621 | Treponema | -2.00 | 1.45 | 1552 | 0.01% | 25.63 | 388 | -1,164 |
| Firmicutes_18412 | Unknown | -2.35 | 1.49 | 697 | 0.00% | 21.41 | 136 | -561 |
| Prevotella_2087 | Prevotella | -2.85 | 1.57 | 1369 | 0.00% | 19.15 | 190 | -1,179 |
| Leptotrichiaceae_8903 | Unknown | -2.63 | 1.32 | 888 | 0.00% | 17.18 | 144 | -744 |

#### Supplementary Table 7. Analysis of *map* gene: frequencies and identities of *Streptococcus* spp.

Results are based on 475 samples, with 37,930,250 reads giving 14,898 *map* gene OTUs

| OUT ID | Identified Species | Abundance | Abundance % | Prevalence | Prevalence % | BLAST | | | |
|---|---|---|---|---|---|---|---|---|---|
| | | | | | | Identity (%) | E Value | Alignment | Tests Seen |
| **OUT10104** | *S. salivarius* | 601,651 | 1.59% | 474 | 99.79 | 99 | 0.00E+00 | 407/412 | 3 |
| **OUT15936** | *S. salivarius* | 424,882 | 1.12% | 465 | 97.89 | 99 | 0.00E+00 | 408/412 | 3 |
| **OUT13812** | *S. salivarius* | 169,574 | 0.45% | 256 | 53.89 | 98 | 0.00E+00 | 403/412 | 2 |
| **OUT10307** | *S. salivarius* | 142,824 | 0.38% | 426 | 89.68 | 99 | 0.00E+00 | 410/412 | 4 |
| **OUT24710** | *S. salivarius* | 96,238 | 0.25% | 457 | 96.21 | 99 | 0.00E+00 | 409/412 | 4 |
| **OUT23481** | *S. salivarius* | 67,211 | 0.18% | 387 | 81.47 | 99 | 0.00E+00 | 408/412 | 3 |
| **OUT21636** | *S. parasanguinis* | 47,040 | 0.12% | 373 | 78.53 | 98 | 0.00E+00 | 405/412 | 3 |
| **OUT11843** | *S. salivarius* | 43,784 | 0.12% | 446 | 93.89 | 98 | 0.00E+00 | 403/412 | 3 |
| **OUT13903** | *S. salivarius* | 43,697 | 0.12% | 366 | 77.05 | 99 | 0.00E+00 | 408/412 | 3 |
| **OUT22893** | *S. salivarius* | 39,182 | 0.10% | 277 | 58.32 | 99 | 0.00E+00 | 406/412 | 2 |
| **OUT10817** | *S. salivarius* | 38,512 | 0.10% | 364 | 76.63 | 99 | 0.00E+00 | 410/412 | 5 |
| **OUT13837** | *S. parasanguinis* | 36,481 | 0.10% | 361 | 76 | 99 | 0.00E+00 | 411/412 | 3 |
| **OUT10706** | *S. salivarius* | 35,934 | 0.09% | 189 | 39.79 | 98 | 0.00E+00 | 402/412 | 3 |
| **OUT10102** | *S. salivarius* | 32,176 | 0.08% | 395 | 83.16 | 99 | 0.00E+00 | 411/412 | 3 |
| **OUT9031** | *S. parasanguinis* | 28,211 | 0.07% | 143 | 30.11 | 94 | 1.00E-175 | 387/411 | 2 |
| **OUT15267** | *S. parasanguinis* | 27,944 | 0.07% | 352 | 74.11 | 94 | 1.00E-175 | 387/411 | 3 |
| **OUT12653** | *S. parasanguinis* | 27,913 | 0.07% | 349 | 73.47 | 95 | 0.00E+00 | 391/411 | 3 |
| **OUT11394** | *S. oralis* | 27,602 | 0.07% | 184 | 38.74 | 85 | 1.00E-110 | 348/411 | 2 |
| **OUT2846** | *S. salivarius* | 26,346 | 0.07% | 359 | 75.58 | 100 | 0.00E+00 | 412/412 | 3 |
| **OUT124** | *S. parasanguinis* | 23,666 | 0.06% | 310 | 65.26 | 94 | 2.00E-172 | 385/411 | 3 |
| **OUT10097** | *S. salivarius* | 23,030 | 0.06% | 451 | 94.95 | 99 | 0.00E+00 | 408/412 | 3 |
| **OUT16404** | *S.* sp. I-G2 | 21,908 | 0.06% | 110 | 23.16 | 88 | 9.00E-132 | 359/408 | 2 |

| | | | | | | | | | |
|---|---|---|---|---|---|---|---|---|---|
| **OUT2013*** | *S. mitis* | 19,246 | 0.05% | 276 | 58.11 | 100 | 1.00E-180 | 348/348 | 4 |
| **OUT10547** | *S. salivarius* | 18,369 | 0.05% | 316 | 66.53 | 99 | 0.00E+00 | 406/412 | 2 |
| **OUT13462** | *S. parasanguinis* | 14,697 | 0.04% | 361 | 76 | 95 | 0.00E+00 | 392/411 | 3 |
| **OUT12401** | *S. salivarius* | 9,055 | 0.02% | 377 | 79.37 | 99 | 0.00E+00 | 406/412 | 2 |
| **OUT1213** | *S. thermophilus* | 6,355 | 0.02% | 265 | 55.79 | 98 | 0.00E+00 | 403/412 | 2 |
| **OUT10860** | *S. parasanguinis* | 5,289 | 0.01% | 61 | 12.84 | 94 | 2.00E-177 | 388/411 | 2 |
| **OUT12371** | *S. salivarius* | 3,705 | 0.01% | 354 | 74.53 | 99 | 0.00E+00 | 406/412 | 2 |
| **OUT13964** | *S. salivarius* | 3,390 | 0.01% | 365 | 76.84 | 99 | 0.00E+00 | 406/412 | 2 |
| **OUT23280** | *S. salivarius* | 3,128 | 0.01% | 254 | 53.47 | 99 | 0.00E+00 | 411/412 | 2 |
| **OUT15822** | *S. salivarius* | 2,997 | 0.01% | 349 | 73.47 | 99 | 0.00E+00 | 411/412 | 2 |
| **OUT14077** | *S. salivarius* | 2,635 | 0.01% | 347 | 73.05 | 99 | 0.00E+00 | 406/412 | 2 |

*OUT2013 potentially *S. pneumoniae* :

| Identity (%) | E Value | Alignment | Tests Seen |
|---|---|---|---|
| 94 | 1.00E-174 | 387/412 | 4 |

Supplementary Table 8. *Streptococcus* spp. affected by smoking

| OTU ID | Identified Species | Abundance % | Prevalence % | DESeq analysis | |
|---|---|---|---|---|---|
| | | | | Fold Change | *P* adjusted |
| **OTU10860** | *S. parasanguinis* | 0.01% | 12.84 | 5.2 | 1.75E-07 |
| **OTU2013** | *S. mitis/pneumoniae* | 0.05% | 58.11 | 3.62 | 4.81E-09 |
| **OTU24710** | *S. salivarius* | 0.25% | 96.21 | 3.03 | 5.59E-15 |
| **OTU1213** | *S. thermophilus* | 0.02% | 55.79 | 2.53 | 7.38E-05 |
| **OTU23280** | *S. salivarius* | 0.01% | 53.47 | 1.82 | 2.59E-04 |
| **OTU12371** | *S. salivarius* | 0.01% | 74.53 | 1.53 | 9.00E-06 |
| **OTU14077** | *S. salivarius* | 0.01% | 73.05 | 1.43 | 2.07E-05 |
| **OTU13964** | *S. salivarius* | 0.01% | 76.84 | 1.34 | 1.00E-04 |
| **OTU12401** | *S. salivarius* | 0.02% | 79.37 | -1.52 | 2.94E-04 |
| **OTU10817** | *S. salivarius* | 0.10% | 76.63 | -1.92 | 3.56E-05 |
| **OTU10706** | *S. salivarius* | 0.09% | 39.79 | -3.82 | 5.96E-08 |
| **OTU11394** | *S. oralis* | 0.07% | 38.74 | -6.6 | 4.38E-17 |
| **OTU16404** | *S.* sp. I-G2 | 0.06% | 23.16 | -8.44 | 2.91E-19 |
